# Early whole-body mutant huntingtin lowering averts changes in proteins and lipids important for synapse function and white matter maintenance in the LacQ140 mouse model

**DOI:** 10.1101/2023.01.26.525697

**Authors:** Kai Shing, Ellen Sapp, Adel Boudi, Sophia Liu, Connor Seeley, Deanna Marchionini, Marian DiFiglia, Kimberly B. Kegel-Gleason

## Abstract

**Background:** Expansion of a triplet repeat tract in exon1 of the HTT gene causes Huntington’s disease (HD). The mutant HTT protein (mHTT) has numerous aberrant interactions with diverse, pleiomorphic effects. No disease modifying treatments exist but lowering mutant huntingtin (mHTT) by gene therapy is a promising approach to treat Huntington’s disease (HD). It is not clear when lowering should be initiated, how much lowering is necessary and for what duration lowering should occur to achieve benefits. Furthermore, the effects of mHTT lowering on brain lipids have not been assessed.

**Methods:** Using a mHtt-inducible mouse model we analyzed whole body mHtt lowering initiated at different ages and sustained for different time-periods. Subcellular fractionation (density gradient ultracentrifugation), protein chemistry (gel filtration, western blot, and capillary electrophoresis immunoassay), liquid chromatography and mass spectrometry of lipids, and bioinformatic approaches were used to test effects of mHTT transcriptional lowering.

**Results:** mHTT protein in cytoplasmic and synaptic compartments of the caudate putamen, which is most affected in HD, was reduced 38-52%. Little or no lowering of mHTT occurred in nuclear and perinuclear regions where aggregates formed at 12 months of age. mHtt transcript repression partially or fully preserved select striatal proteins (SCN4B, PDE10A). Total lipids in striatum were reduced in LacQ140 mice at 9 months and preserved by early partial mHtt lowering. The reduction in total lipids was due in part to reductions in subclasses of ceramide (Cer), sphingomyelin (SM), and monogalactosyldiacylglycerol (MGDG), which are known to be important for white matter structure and function. Lipid subclasses phosphatidylinositol (PI), phosphatidylserine (PS), and bismethyl phosphatidic acid (BisMePA) were also changed in LacQ140 mice. Levels of all subclasses other than ceramide were preserved by early mHtt lowering. Pathway enrichment analysis of RNAseq data imply a transcriptional mechanism is responsible in part for changes in myelin lipids, and some but not all changes can be rescued by mHTT lowering.

**Conclusions:** Our findings suggest that early and sustained reduction in mHtt can prevent changes in levels of select striatal proteins and most lipids but a misfolded, degradation-resistant form of mHTT hampers some benefits in the long term.

## BACKGROUND

Huntington’s disease (HD) is a heritable neurodegenerative disease caused by a CAG expansion in the huntingtin (*HTT*) gene. The protein product, huntingtin (HTT), is ubiquitously expressed but enriched in neurons ^1,2^ and has been implicated in numerous molecular functions including vesicle trafficking, autophagy, transcription and DNA repair (Reviewed by ^3–5^). HTT has also been shown to have essential functions during development ^6–11^. The CAG-repeat expansion encodes a polyglutamine expansion in HTT, which causes protein accumulation and aggregation and has pleiomorphic effects that contribute to HD pathology ranging from mitochondrial dysfunction, transcriptional defects, cholesterol mishandling, altered palmitoylation, and metabolic changes from altered signaling in the hypothalamus ^12,13^.

*HTT* transcription lowering strategies have become central to HD translational studies (reviewed by ^14^). Interventional trials have targeted total *HTT* (that is, normal and mutant *HTT*) using antisense oligonucleotides (ASOs) (reviewed by ^15^) or *HTT* pre-mRNA using an oral drug, branaplam (LMI070) ^16^. In animal models (mice, sheep, and nonhuman primates), additional strategies of gene therapy using transcriptional repressors to target expression from the mutant allele ^17^ and modified interfering RNAs (RNAi) ^18^ or AAV expressing microRNAs (miRNAs) or short hairpin RNAs (shRNAs) ^19,20^ to target RNA levels are actively being pursued ^17,21–28^.

Although lowering total *HTT* in humans was hoped to be generally safe ^29^, insufficient evidence currently exists to make this conclusion. There is support for selective lowering of m*HTT* based on data in mice that show loss of wild type (WT) *HTT* may affect neuronal function including synaptic connectivity ^6,8,30,31^.

Proteins known to change with HD progression may serve as useful readouts for investigating the efficacy of HTT lowering. For instance, DARPP32 is enriched in striatal projection neurons and is progressively reduced in HD patient postmortem brain and in mouse HD models ^32^. The cAMP phosphodiesterase PDE10A is reduced early and sustainably in HD striatum measured both by western blot and mass spectrometry ^17,33,34^ or ligand binding ^17,35^. Microarray analysis in neurons derived from human stem cells ^36^, mass spectrometry and western blot analyses in striatal synaptosomes ^33^ and immunofluorescence (IF) studies in mice ^37^ showed SCN4B is lowered in HD models. ATP5A protein levels are altered in numerous mass spectrometry studies ^33,38–40^.

The intracellular location of HTT positions it to affect lipids in membranes. HTT normally associates with membranes where it interacts directly with lipid bilayers ^41^. Lipids comprise ∼50% of the total dry weight in brain and have diverse cellular roles as membrane structural components, signaling molecules, and sources of energy ^42^. Disruptions to lipid composition/metabolism have been linked to disease ^43^; thus, their study may provide insight into disease mechanism or lead to discovery of new biomarkers. We and others have described alterations to lipid levels in HD mouse models ^44–48^ and post-mortem HD brain tissue ^49–51^. Specific glycerophospholipids showed differential associations with mHTT ^52^ and mHTT can alter the biophysical properties of lipid bilayers ^41,53^ underscoring the potential for direct consequences of the presence of mHTT on cellular lipids.

The effects of lowering total *Htt* or m*Htt* alone on striatal proteins and behavioral and psychiatric measures have been investigated in HD mouse models after delivery of reagents to the striatum or lateral ventricle ^17,54^. Biomarkers (mHTT levels and aggregation) that are responsive to total or m*Htt* lowering have been investigated in HD mouse models ^35,55,56^. However, the impact of lowering m*Htt* on lipids has not been examined. Here we used the inducible HD knock-in mouse model, LacQ140, in which the expression of m*Htt* alone is regulated by adding or withdrawing the lactose analog isopropyl b-D-1-thiogalactopyranoside (IPTG) in their drinking water ^57–60^, to study the effects of m*Htt* lowering for different time periods on proteins known to be affected in HD models and using a survey of lipids. We also tracked changes in levels of soluble and aggregated mHTT protein in different subcellular compartments. The results show that early and sustained reduction in m*Htt* in this HD mouse model can delay select protein changes and prevent numerous lipid derangements. Using a systems biology approach, we integrated our lipidomic findings with a previously generated transcriptomic dataset, identifying dysregulation of pathways relating to myelin and white matter including genes directly involved in biosynthetic pathways for lipids we found changed in LacQ140 mice. An SDS-soluble misfolded form of mHTT persists, however, and benefits eventually dwindled.

## MATERIALS AND METHODS

### Animals

The *LacO*/*LacIR*-regulatable HD mouse model (LacQ140) was generated by crossing the *Htt^LacQ140/+^*mouse to the *Tg^ACTB-lacI*Scrb^* mouse as previously described ^58,59^. The default state of the LacQ140 mouse is global repression of m*Htt* due to *Lac* Repressor binding to the *Lac* operators. The continuous administration of IPTG starting from E5 interrupts the binding between the *Lac* repressor and operators, resulting in a de-repressed state, and maximal expression of m*Htt* in LacQ140. All WT mice were Htt^LacO+/+^; b-actin-LacI^R^ tg. Mice were provided with enrichment (envirodry, play tunnels, Bed-o’cobs and plastic bones) and housed uniform for genotype, gender, and treatment. Mice were fed *ad libitum*. The lactose analog IPTG was provided in drinking water (at 10mM) which de-represses the *LacQ140* allele and keeps normal m*Htt* expression. During embryonic development, m*Htt* expression levels were maintained at normal levels by administering IPTG to pregnant dams starting at embryonic day 5 (E5). IPTG was administered never (*mHtt* repressed), always (*mHtt* always expressed) or withdrawn at 2 or 8 months (*mHtt* expressed normally then repressed at 2 or 8 months). The CAG repeat length range in *Htt^LacO-Q140/+^*mice was 143-157 with average of 148 and median of 148 CAG.

### Sample preparation

The striatum from one hemisphere for each mouse was homogenized in 750 µl 10mM HEPES pH7.2, 250mM sucrose, 1uM EDTA + protease inhibitor tablet (Roche Diagnostics GmbH, Mannheim, Germany) + 1mM NaF + 1mM Na3VO4. A 150 µl aliquot of this crude homogenate was removed and protein concentration was determined using the Bradford method (BioRad, Hercules, CA). Subcellular fractionation by density gradient ultracentrifugation using Optiprep was performed on remaining 600 µl sample for the 6- and 12-month-old mice as previously described ^47^.

### Capillary immunoassay

Equal amounts of protein from the crude homogenates were analyzed using the automated simple western system, Wes (ProteinSimple, Bio-Techne, San Jose, CA), which utilizes a capillary-based immunoassay. The protocol described in the manual was followed to detect HTT, GFAP and DARPP32 using 0.6 µg of sample. Quantitative analysis of protein levels is done automatically using the Compass for Simple Western Software (ProteinSimple) on electropherogram data. The peak area (using automatic “dropped line” option in software) of each protein of interest was normalized to the peak area of the vinculin loading control. Figures show protein bands similar to traditional western blots using “lane view” option in the Compass software to create a blot-like image from the electropherogram data.

### Western blot analysis

Equal amounts of protein from the crude homogenates were analyzed by western blot for levels of HTT and other proteins of interest as previously described ^33^. Briefly, 10 µg of protein were separated by SDS-PAGE, transferred to nitrocellulose, and probed with primary antibody overnight. Nitrocellulose membranes were cut into horizonal strips to maximize the number of antibodies that could be probed on one blot. Peroxidase labeled secondary antibodies were used with the SuperSignal West Pico Chemiluminescent substrate (ThermoScientific, Rockford, IL, #34580) and digital images were captured with a CCD camera (AlphaInnotech, Bayern, Germany). For western blot analysis of subcellular fractions, equal volumes of each sample (15 µl) were separated by SDS-PAGE. Pixel intensity quantification of the western blot signals on the digital images was determined using ImageJ software (NIH) by manually circling each band and multiplying the area by the average signal intensity. The total signal intensity was normalized to vinculin or GAPDH loading controls.

### Antibodies

The following antibodies and dilutions were used in this study: Anti-HTT Ab1 (aa1-17,^1^) 1:50 for capillary immunoassay and 1:2000 for western blot; anti-HTT EPR5526 (Abcam, Waltham, MA, ab109115, 1:2000 for western blot); anti-polyQ MW1 (MilliporeSigma, Burlington, MA, MABN2427, 1:50 for capillary immunoassay); anti-polyQ PHP3 (generous gift from Dr. Ali Khoshnan, 1:2000 for western blot); Anti-PDE10A (Abcam, Waltham, MA, #ab177933, 1:2000 for western blot); Anti-DARPP32 (Abcam, #ab40801, 1:2000 for capillary immunoassay); Anti-GFAP (MilliporeSigma, Burlington, MA, AB5804, 1:3000 for capillary immunoassay); Anti-GAPDH (MilliporeSigma, Burlington, MA, #MAB374, 1:10000 for western blot); Anti-Sodium channel subunit beta-4 (Abcam, Waltham, MA, #ab80539, 1:500 for western blot); Anti-vinculin (Sigma, St. Louis, MO, #V9131, 1:5000 for capillary immunoassay, 1:2000 for western blot); Anti-ATP5A (Abcam, Waltham, MA, #ab14748, 1:2000 for western blot); Anti-HTT MW8 (University of Iowa Developmental Studies Hybridoma Bank, 1:1000 for filter trap); Anti-HTT S830 (generous gift from Dr. Gillian Bates, ^61^ 1:8000), HDAC1 (Abcam, Waltham, MA, ab32369-7, 1:4000).

### Filter trap assay

Based on protocol described in ^62,63^, equal protein amounts for each sample (40 µg) were brought up to 50 µl volume with PBS and 50 µl 4% SDS in PBS was added to each sample to make final concentration 2% SDS. A cellulose acetate membrane was wet in 2% SDS/PBS and placed in dot blot apparatus. The 100 µl samples were added to each well and pulled through the membrane with a vacuum then washed 3 times with 200 µl 2% SDS/PBS. The membrane was removed from the apparatus, washed in Tris buffered saline + 0.1% Tween-20 (TBST) then processed for western blot using MW8 or S830 antibodies. The total signal intensity of each dot was measured in ImageJ by circling the entire dot and multiplying the area by the average signal intensity minus the background signal from an empty dot.

### Statistical analysis

One-way ANOVA with Tukey’s multiple comparison test was performed to determine significance between groups. Asterisks on graphs show p values and are described in the figure legends.

### Lipid extraction

Lipids were extracted using methyl tert-butyl ether (MTBE) as previously described and analyzed using ion switching and molecular assignment as previously described ^47,64,65^. Each age group was processed together. Crude homogenates (100 µl) of dissected mouse striatum were transferred into 20 ml glass scintillation vials. 750 µl of HPLC grade methanol was added to each sample, then vials were vortexed. 2.5 ml of MTBE was then added to each sample and incubated on a platform shaker for 1 hour. After incubation, 625 µl of water was added to induce polar and non-polar phase separation. The non-polar lipid containing (upper) phase was collected into a new vial, and the polar (lower) phase was subsequently re-extracted with 1 ml of MTBE/methanol/water (10/3/2.5, v/v/v). Following re-extraction, the lipid containing phases were combined and allowed to dry on a platform shaker, then further dried with nitrogen gas. Extracted lipids were hand delivered to the Beth Israel Deaconess Medical Center Mass Spectrometry Core Facility.

### Lipid Annotation

Data for each timepoint was classified by LIPID MAPS category^66^: glycerophospholipids, glycerolipids, sphingolipids, sterol lipids, fatty acyls, and prenol lipids were detected. Each category contains distinct subclasses as annotated below. *<u>Glycerophospholipids:</u>*Phosphatidylcholine (PC), Phosphatidylethanolamine (PE), Phosphatidylserine (PS), Phosphatidylinositol (PI), Methylphosphocholine (MePC), Phosphatidic acid (PA), Bis-methyl phosphatidic acid (BisMePA), Dimethyl phosphatidylethanolamine (dMePE), Phosphatidylgylcerol (PG), Bis-methylphosphatidylserine (BisMePS), Bis-methyl phosphatidyl ethanolamine (BisMePE), Cardiolipin (CL), Phosphatidylethanol (PEt), Biotinyl-phosphoethanolamine (BiotinylPE), Phosphatidylmethanol (PMe), Phosphatidylinositol-bisphosphate (PIP2), Phosphatidylinositol-monophosphate (PIP), Lysophosphatidylcholine (LPC), Lysophosphatidylethanolamine (LPE), Lysophosphatidylserine (LPS), Lysophosphatidylinositol (LPI), Lysophosphosphatidylgylcerol (LPG), Lysodimethyl phosphatidyl ethanolamine (LdMePE). <u>*Glycerolipids:*</u> Triglyceride (TG), Monogalactosyldiacylglycerol (MGDG), Monogalactosylmonoacylglycerol (MGMG), Diglyceride (DG), Sulfoquinovosylmonoacylglycerol (SQMG), Sulfoquinovosyldiacylglycerol (SQDG). *<u>Sphingolipids:</u>* Hexosylceramides (Hex1Cer), Simple Glc series (CerG1), Sphingomyelin (SM), Ceramide (Cer), Ceramide phosphate (CerP), Sulfatide (ST), Sphingoid base (So), Sphingomyelin phytosphingosine (phSM), Simple Glc series (CerG2GNAc1), Ceramide phosphorylethanolamine (CerPE), Sphingosine (SPH), Dihexosylceramides (Hex2Cer). *<u>Sterol lipids</u>*: Cholesterol ester (ChE), Zymosterol (ZyE). *<u>Fatty acyls:</u>* Fatty acid (FA), Acyl Carnitine (AcCa). *<u>Prenol lipids:</u>* Coenzyme (Co).

Individual lipid species were annotated according to sum composition of carbons and double bonds in the format Lipid Subclass (total number of carbons: total number of double bonds). If distinct fatty acid chains could be identified, they were annotated separated by an underscore (ex. PC 32:1, or PC (16:0_16:1). Using this approach, we cannot determine the *sn*-1 or *sn*-2 positions of the fatty acid chains. Lipid species within the sphingolipid category contain prefixes ‘d’ or ‘t’ to denote di-hydroxy or tri-hydroxy bases. For example, SM(d18:1_23:0) contains 2 hydroxyl groups. The Hex1Cer subclass is comprised of both glucosylceramide (GlcCer) and galactosylceramide (GalCer); the orientation of one of the hydroxyl groups in Glc differs from in Gal, and thus cannot be resolved by these methods ^67^. Plasmanyl lipid species (ether linked) are annotated by ‘e’ and plasmenyl/plasmalogen (vinyl ether linked) lipid species are annotated by ‘p’ (ex. PC (36:5e) or PE (16:0p_20:4) ^68^.

### Lipidomics statistics and data visualization

Heatmap and hierarchical clustering in Figure 4 were generated using Morpheus from the Broad Institute (Cambridge, MA, https://software.broadinstitute.org/morpheus). Hierarchical clustering was performed across all rows (lipid subclasses) using the one minus Pearson correlation distance metric. Rows determined to be the most similar are merged first to produce clusters, working iteratively to merge rows into clusters. The dendrogram displays the order of clustering with the most similar rows shown in closest proximity. Lipid expression values are assigned to colors based on the minimum (blue, low relative expression) and maximum (red, high relative expression) values for each independent row. Heatmap in Figures 4 & 5 was generated in R using the ComplexHeatmap package v2.16.0 ^69^. Each column represents data from one animal. Statistical significance was determined by one-way analysis of variance (ANOVA) with Tukey’s multiple comparison test between treatment groups for lipid subclasses (6mo: N=36, 9mo: N=24, 12mo: N=29) or lipid species (6mo: N=800, 9mo: N=632, 12mo: N=735). The two-stage linear step-up procedure of Benjamini, Krieger, and Yekutieli was applied to ANOVA p-values to control the false discovery rate (FDR) with significance accepted at q<0.05 (Q=5%).

### RNA-sequencing analysis

The raw counts matrix derived from striatal samples in the LacQ140 mouse model was accessed from Gene Expression Omnibus (GEO GSE156236). The 6-month cohort included n=10 mice per group (WT, LacQ140, LacQ140_2M, LacQ140_A) and the 12-month cohort included n=10 WT, n=9 LacQ140, n=9 LacQ140_8M, n=10 LacQ140_2M, and n=10 LacQ140_A. Analysis was performed using R v4.3.0 in the RStudio IDE (2023.03.0+386). Transcripts with fewer than 10 total counts were removed from the analysis. Differential expression analysis was performed using DESeq2 v1.40.1 ^70^. P-values were computed with the Wald test and were corrected for multiple testing using the Benjamini and Hochberg procedure ^71^. Genes with a fold change greater than 20% in either direction and with an adjusted p-value < 0.05 were considered differentially expressed. Functional enrichment analysis was conducted using the ClusterProfiler package v4.8.1. Over-representation analysis was performed for upregulated or downregulated DEGs with the enrichGO function (org.Mm.eg.db v3.17.0). Gene set enrichment analysis was performed with the gseGO function using a ranked list of genes (Wald statistic) for each comparison (LacQ140 vs WT, LacQ140_8M vs WT, LacQ140_2M vs WT, and LacQ140_A vs WT) at 6 and 12 months. Heatmaps were plotted using ComplexHeatmap package v2.16.0 with rows representing individual animals and rows representing genes.

## RESULTS

### Time course of mHTT protein lowering with regulated transcriptional repression using LacQ140 mouse striatum

We used the inducible HD knock-in mouse model, LacQ140, in which the expression of m*Htt* throughout the body was regulated by adding or withdrawing the lactose analog isopropyl b-D-1-thiogalactopyranoside (IPTG) in drinking water using an established *Lac* operator/repressor system (**Fig. 1a**) ^59,60^. We used the model to lower m*Htt* from conception, starting at 2 or 8 months of age. To control for the effects of IPTG, WT mice also received IPTG treatment over their lifetime. The striatum was examined at 6, 9, and 12 months of age (**Fig. 1b**). Antibodies against varied epitopes within HTT detect different pools of normal and mutant HTT by immunofluorescence and immunohistochemistry ^1,72–75^. Numerous conformations (over 100 distinct classes) of purified HTT and mHTT have been identified by electron microscopy in vitro ^76^. Polyglutamine (polyQ) expansion induces confirmational changes particular to mHTT detectible even after SDS-PAGE and/or western blot with several antibodies to regions in the poly-Q region (MW1 ^77,78^ and 3B5H10 ^79^) and in the adjacent polyproline region (PHP1-4 ^80^ and S830 ^81^). To assess lowering, full length HTT and mHTT proteins were detected by capillary immunoassay using anti-HTT antibody Ab1 targeting aa 1-17 and anti-polyQ antibody MW1 (**Fig. 2**), and by western blot using anti-HTT antibody EPR5526, and anti-mHTT antibody PHP3, which recognizes an altered conformer of polyproline in mHTT ^80^ (**Fig. S1**).

**Figure 1.**
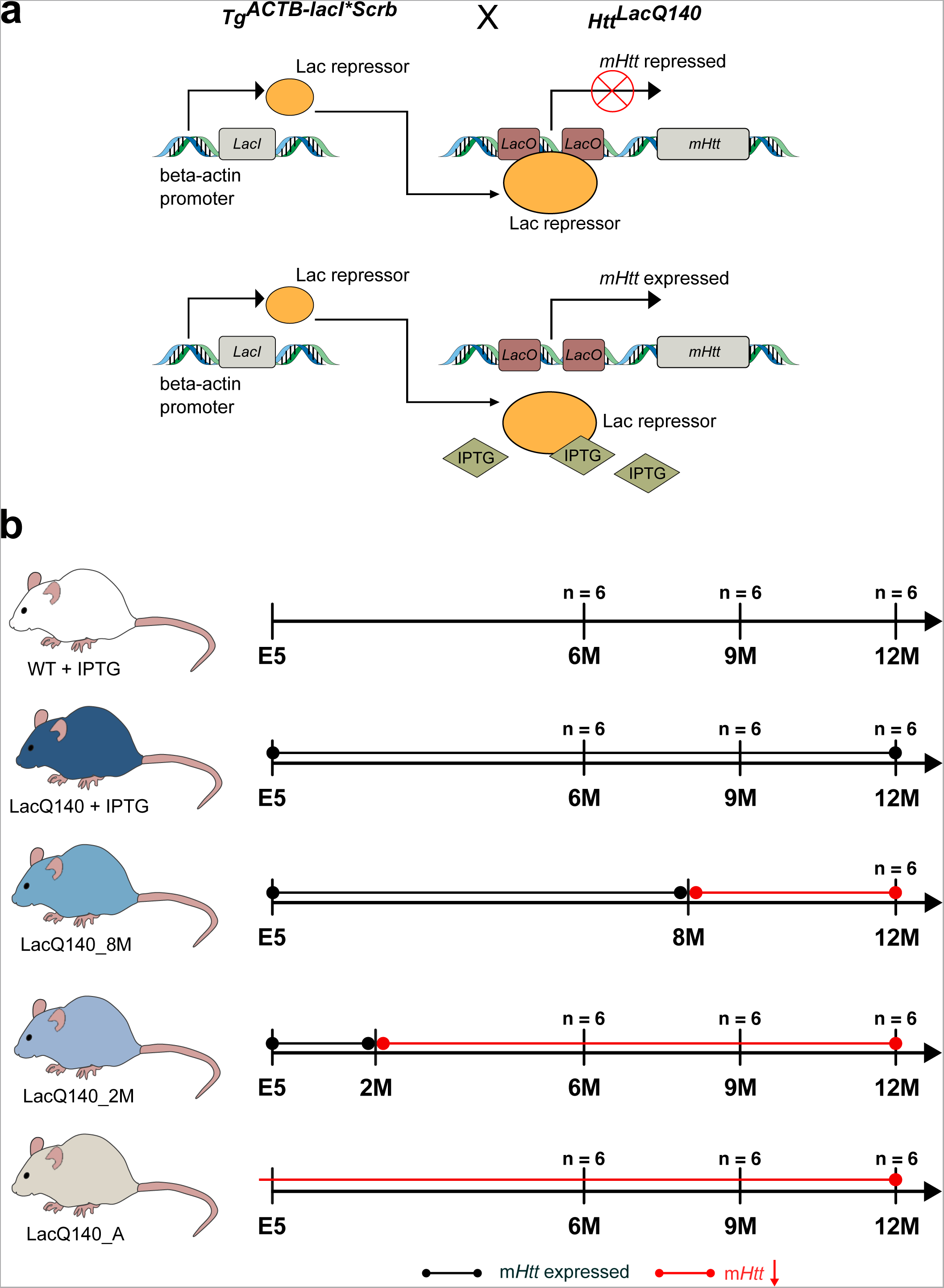
Generation of LacQ140 mice and treatment paradigm (a) The *LacO*/*LacIR*-regulatable HD mouse model (LacQ140) was generated by crossing the *Htt^LacQ140/+^* mouse to the *Tg^ACTB-lacI*Scrb^* mouse ^58^ as previously described ^59^. The default state of the LacQ140 mouse is global repression of m*Htt* due to *Lac* Repressor binding to the *Lac* operators. Administration of IPTG starting from embryonic day 5 (E5) interrupts the binding between the *Lac* repressor and operators, resulting in a de-repressed state, and maximal expression of m*Htt* in LacQ140. All WT mice were Htt^LacO+/+^; b-actin-LacI^R^ tg. **(b)** Mice were fed *ad libitum*; the lactose analog IPTG was provided in drinking water (at 10mM) which de-represses the *LacQ140* allele and keeps normal m*Htt* expression. During embryonic development, m*Htt* expression levels were maintained at normal levels by administering IPTG to pregnant dams starting at embryonic day 5 (E5). IPTG was continuously administered to WT mice. IPTG was administered always (m*Htt* always expressed, LacQ140), withdrawn at 8 months (m*Htt* repressed beginning at 8 months, LacQ140_8M), withdrawn 2 at months (m*Htt* repressed beginning at 2 months, LacQ140_2M), or never administered (m*Htt* always repressed, LacQ140_A). Tissue for each group (except LacQ140_8M) was collected at 6, 9, and 12 months of age.

**Figure 2.**
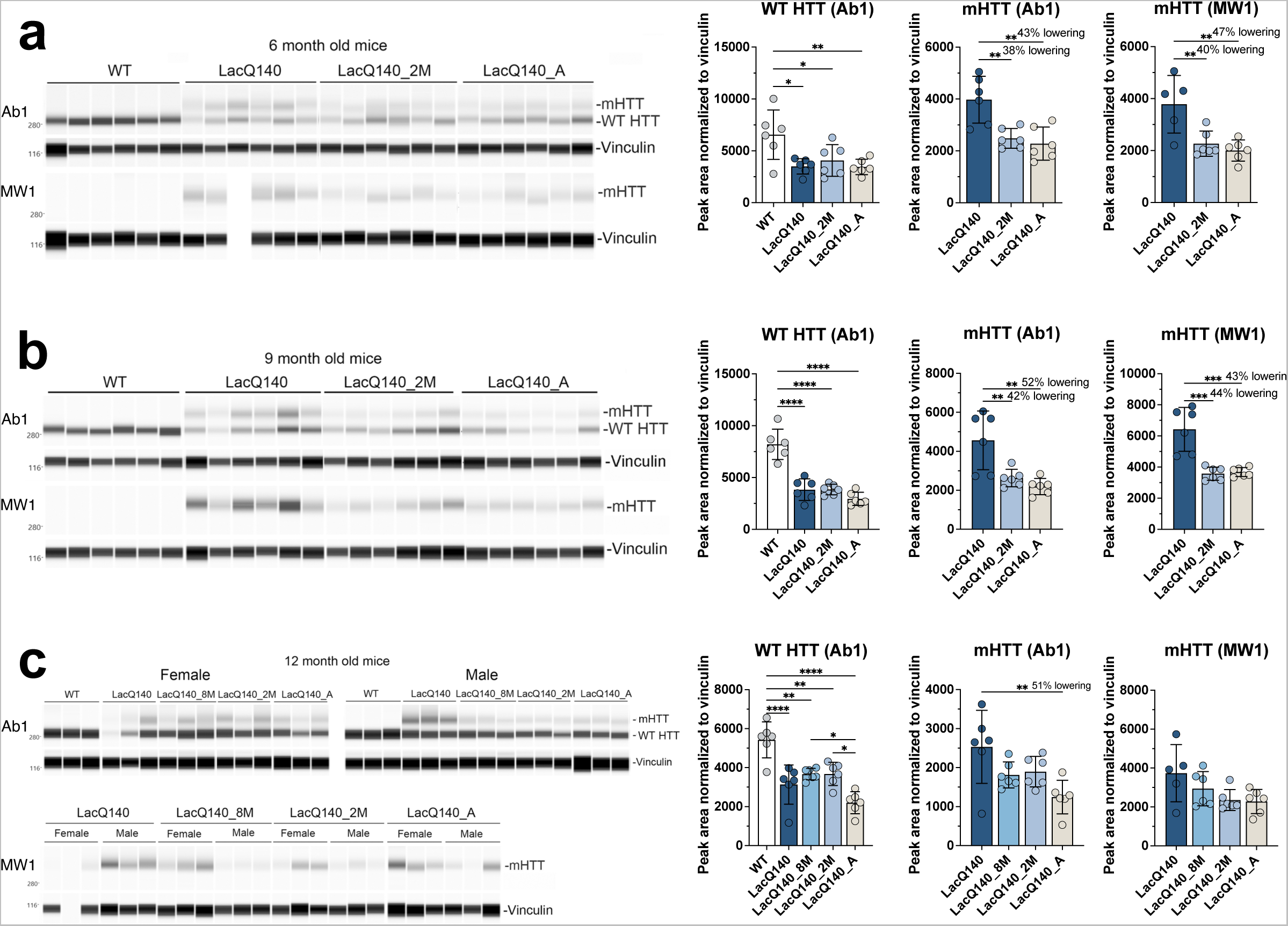
Analysis of mHTT protein levels in crude homogenates of 6-, 9- and 12-months old mice. HTT levels were analyzed by capillary immunoassay on equal amounts of protein (0.6 µg) using anti-HTT antibody Ab1 and anti-polyQ antibody MW1 (**a**). Peak area analysis performed using Compass software in 6-month-old mice shows a significant decrease in WT HTT as detected with Ab1 in all LacQ140 mice compared to WT mice (F(3, 20) = 5.674, **P=0.0056, One-way ANOVA with Tukey’s multiple comparison test, n=6). mHTT levels are significantly lower in LacQ140_2M and LacQ140_A as detected with both Ab1 and MW1 compared to LacQ140 (**a**, Ab1: F(2, 15) = 11.25, **P=0.0010, -38% and -43% respectively; MW1: F(2, 14) = 9.879, **P=0.0021, -40% and -47% respectively). Peak area analysis in 9-month-old mice shows a significant decrease in WT HTT as detected with Ab1 in all LacQ140 mice compared to WT mice (F(3, 20) = 34.67, ****P<0.0001, One-way ANOVA with Tukey’s multiple comparison test, n=6). mHTT levels are significantly lower in LacQ140_2M and LacQ140_A, as detected with both Ab1 and MW1, compared to LacQ140 (**b**, Ab1: F(2, 15) = 10.82, **P=0.0012, -42% and -52% respectively; MW1: F(2, 15) = 20.82, ****P<0.0001, -44% and -43% respectively). Peak area analysis in 12-month-old mice shows a significant decrease in WT HTT as detected with Ab1 in all LacQ140 mice compared to WT mice (F(4, 25) = 15.81, ****P<0.0001, One-way ANOVA with Tukey’s multiple comparison test, n=6). WT HTT was significantly lower in LacQ140_A compared to LacQ140_8M and LacQ140_2M mice. mHTT levels are significantly lower in LacQ140_A mice, as detected with Ab1, compared to LacQ140 (**c**, Ab1: F(3, 20) = 5.017, **P=0.0094, -51%, One-way ANOVA with Tukey’s multiple comparison test, n=6). Asterisks on graphs represent Tukey’s multiple comparison test, n=6 mice per group (*p<0.05, **p<0.01, ***p<0.001, ****p<0.0001).

Overall, results showed lowering of 35-52% of mHTT detected with Ab1, MW1 and PHP3 in 6- and 9-month mice with transcriptional repression (**Fig. 2a**, **Fig. S1a**) and (**Fig. 2b**, **Fig. S1b**). Although EPR5526 recognizes both WT and mHTT, no significant lowering of mHTT protein was measured with mHtt gene repression using this antibody in 6-month mice. In the 12-month mice, mHTT was significantly reduced in LacQ140_A (51%) mice compared to LacQ140 mice (**Fig. 2c**). However, this was observed only in the capillary immunoassay using antibody Ab1 but not with antibodies MW1, EPR5526, or PHP3 **(Fig. S1c)**. m*Htt* repression in the LacQ140_2M and LacQ140_8M mice yielded no significant reduction of mHTT protein levels compared to LacQ140 mice using any antibody. These results show that systemic regulated repression of m*Htt* transcription in LacQ140 mice results in partial lowering of mHTT protein levels in the striatum (38-52%), but a soluble form of full-length mHTT remains as mice age.

### Effects of m*Htt* lowering on its distribution in different subcellular cytoplasmic compartments

We next looked at the effects of m*Htt* repression on its protein levels in different subcellular compartments. Density gradient fractionation and ultracentrifugation for subcellular fractionation of cytoplasmic components was performed as shown in **Fig. S2a**. The schematic in **Fig. S2b** indicates representative proteins that are enriched in different cytoplasmic compartments. In 6-month-old mice, the mHTT/WT HTT ratio was significantly lower in LacQ140_2M mice in fractions 13 and 14 compared to LacQ140 mice (**Fig. S2c and d**). Similar results were observed in 12-month-old mice, where the mHTT/WT HTT ratio in fractions 1, 3, 13, and 14 in LacQ140 mice was significantly lower with m*Htt* repression for different periods of time compared to LacQ140 mice (**Fig. S2e and f**). There was no change in the distribution of HTT and mHTT in the fractions between groups in 6-month-old mice **(Fig. S3a and b)** or in 12-month-old mice **(Fig. S3c and d)**. Altogether, these results show that mHTT is lowered in cytoplasmic fractions through 12 months.

### Effects of m*Htt* lowering on its distribution in crude nuclear fractions

In 12-month-old LacQ140 mice, repressing m*Htt* transcription only partially reduced levels of mHTT protein in crude homogenates (**Fig. 2c**) even though mHTT was efficiently lowered in the sub cytoplasmic compartments contained in the S1 fraction (**Fig. S2f**). We speculated that mHTT may reside in other compartments where it is more resistant to removal by transcript repression. To address this idea, we examined P1 fractions which contain nuclei, ER, large perinuclear structures such as the recycling compartment, some mitochondria and autophagosomes ^82^. HTT was detected with antibody Ab1 in WT mice (2 alleles worth) and LacQ140 mice (1 allele worth) and mHTT (1 allele worth) in LacQ140 mice at both 6 and 12 months (**Fig. 3a and b**). Significant lowering of mHTT protein was observed in 6-month-old LacQ140_A but not LacQ140_2M mice using antibody Ab1 (**Fig. 3a**). In 12-month mice, repression of the m*Htt* allele at any age or duration failed to lower mHTT protein levels in the P1 fraction (**Fig. 3b**).

**Figure 3.**
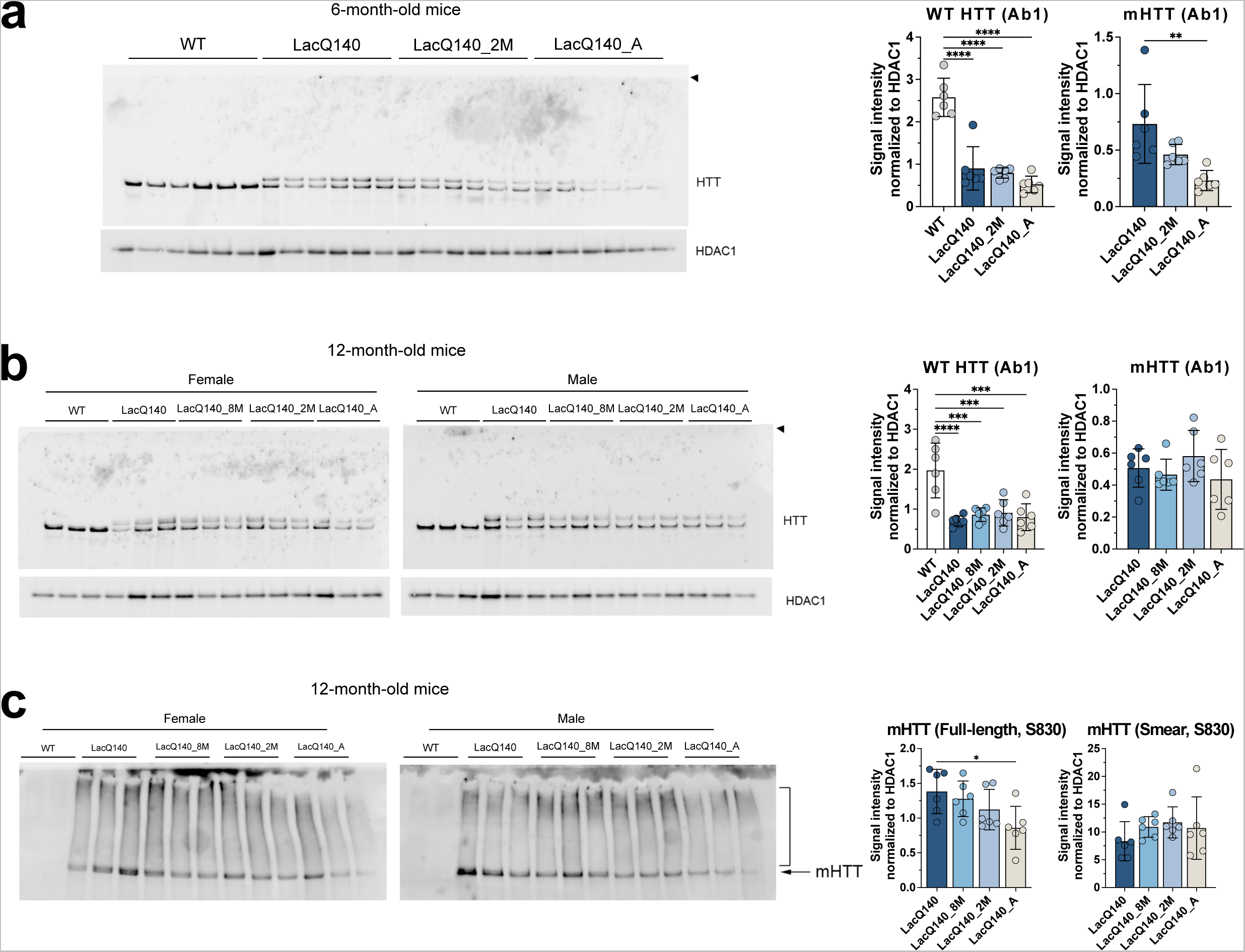
HTT protein levels in P1 fractions. Equal protein (10 µg) from P1 fractions from 6-month-old (**a**) and 12-month-old (**b**) LacQ140 and WT mice were analyzed by western blot for HTT levels with anti-HTT Ab1. No aggregated protein was observed at the top of the gel (arrowhead). Total pixel intensity quantification for each band was measured using ImageJ software and normalized to HDAC1 signal. There was a significant decrease in WT HTT signal in all the treatment conditions for LacQ140 mice compared to WT mice in both **(a)** 6-month-old mice (F(3, 20) = 40.34, ****P<0.0001, One-way ANOVA with Tukey’s multiple comparison test, n=6) and **(b)** 12-month-old mice (F(4, 25) = 11.01, ****P<0.0001, One-way ANOVA with Tukey’s multiple comparison test, n=6). There were significantly lower levels of mHTT in the 6-month-old LacQ140_A mice compared to LacQ140 (**a,** F(2, 15) = 8.233, **P=0.0039, One-way ANOVA with Tukey’s multiple comparison test, n=6) but no changes in mHTT levels were detected in the **(b)** 12-month-old LacQ140 mice (F(3, 20) = 1.137, P=0.3583, n.s., One-way ANOVA). Equal protein (10 µg) from P1 fractions from 12-month-old LacQ140 and WT mice were analyzed by western blot for HTT levels with anti-HTT S830 (**c**). The S830 antibody detected a smear of HTT signal (bracket) as well as full-length mHTT (arrow). There were significantly lower levels of full length mHTT in the 12- month-old LacQ140_A mice compared to LacQ140 (F(3, 20) = 3.548, *P=0.0330, One-way ANOVA with Tukey’s multiple comparison test, n=6) and no changes detected in the HTT smear in all LacQ140 mice (F(3, 20) = 0.9281, P=0.4453, n.s., One-way ANOVA). Asterisks on graphs represent Tukey’s multiple comparison test, n=6 mice per group (*p<0.05, **p<0.01, ***p<0.001, ****p<0.0001).

We queried whether forms of mHTT with altered migration with SDS-PAGE could be detected in the P1 fractions using antibody S830, which has been reported by us and others to detect a smear using SDS-PAGE and western blot ^61,83^. At 12-months, HD mice showed an S830-positive smear above the HTT/mHTT bands which was not lowered in the LacQ140_8M, LacQ140_2M, or LacQ140_A mice (**Fig. 3c**). Filter trap assay showed lowering of an SDS- insoluble aggregated species (recognized by S830) in both crude homogenate and P1 fractions; slightly less lowering of aggregated mHTT occurred in the P1 fraction when transcript repression was initiated at 2 or 8 months (**Fig. S4a and b**). Altogether, these results show that a soluble species of mHTT in the P1 fraction is more resistant to lowering after transcriptional repression.

### Effects of m*Htt* lowering on levels of GFAP, DARPP32, SCN4B, PDE10A, and ATP5A

Prior studies in different mouse models of HD have shown that levels of some neuronal proteins are altered in the mouse striatum, namely DARPP32, PDE10A, SCN4B, and ATP5A ^32–34,37–40,84^. To assess levels of these proteins and that of the astrocyte protein, glial acidic fibrillary protein (GFAP), in LacQ140 mice without and with m*Htt* gene repression, crude homogenates from 6, 9, and 12-month-old mice were analyzed by capillary immunoassay or western blot.

In agreement with previous studies in HD mouse models ^17,33,34^ striatum from the LacQ140 exhibited a significant reduction in PDE10A at all ages examined. Early m*Htt* lowering starting at 2 months of age statistically preserved PDE10A levels at 6 and 9 months (**Fig. S5a and b**), but the effect was lost by 12 months of age when transcript repression was initiated at 2 or 8 months (**Fig. S5c**). Transcript repression initiated embryonically (LacQ140_A) and examined at 6 and 12 months preserved PDE10A levels, although no protection was observed at 9 months. SCN4B expression was reduced in the striatum of LacQ140 at 6 and 9 months of age and levels were preserved with early m*Htt* lowering (**Fig. S5a and b**). DARPP32 levels were significantly lower in LacQ140 compared to WT mice at 9 months but not at 6 or 12 months, and there were no differences in levels of GFAP and ATP5A between WT and LacQ140 at any age examined (**Fig. S6a-c**).

### Effects of m*Htt* lowering on lipids detected by mass spectrometry

We surveyed for lipid changes in LacQ140 striatum compared to WT and the effects of lowering m*Htt* using liquid chromatography and mass spectrometry (LC-MS) as previously described ^47^. For each age, lipids were extracted from crude homogenates of striatum from each treatment group/genotype and analyzed as a set. The total lipids per group were compared. Our MS intensity measurements were relative measurements so only samples processed together can be compared (i.e., by age group). The sum of lipids for each genotype and/or treatment group were reported as a proportion of WT within each age group (**Fig. 4a-c**). No changes in total lipid were observed at 6 months or 12 months (**Fig. 4a and c**). However, at 9 months LacQ140 mice had significantly lower levels of total lipids compared to WT or to LacQ140_A mice (**Fig. 4b**). A heat map and hierarchical clustering of lipid changes by subclass at 9 months revealed two major groups where-in subclasses moved in the same direction even if all weren’t statistically significant (**Fig. 4d**). The top cluster delimited in blue shows subclasses that decreased in LacQ140 mice compared to WT and were corrected by lowering. In contrast, the cluster marked in red shows subclasses that were increased in LacQ140 mice compared to WT and were improved by m*Htt* lowering. Other subclasses marked in black did not change. Summaries of lipid subclasses and number of species detected for each age group are in **Table 1**, and an overview of changes in individual species across ages is shown in **Table 2**. Data and graphs for 6-month-old mice subclasses and species can be found in **Fig. S7 & 8** and **Additional File 1**; 9-month-old mice in **Fig. S9 & 10** and **Additional File 1**; and 12-month-old mice in **Fig. S11 & 12** and **Additional File 1**.

**Figure 4.**
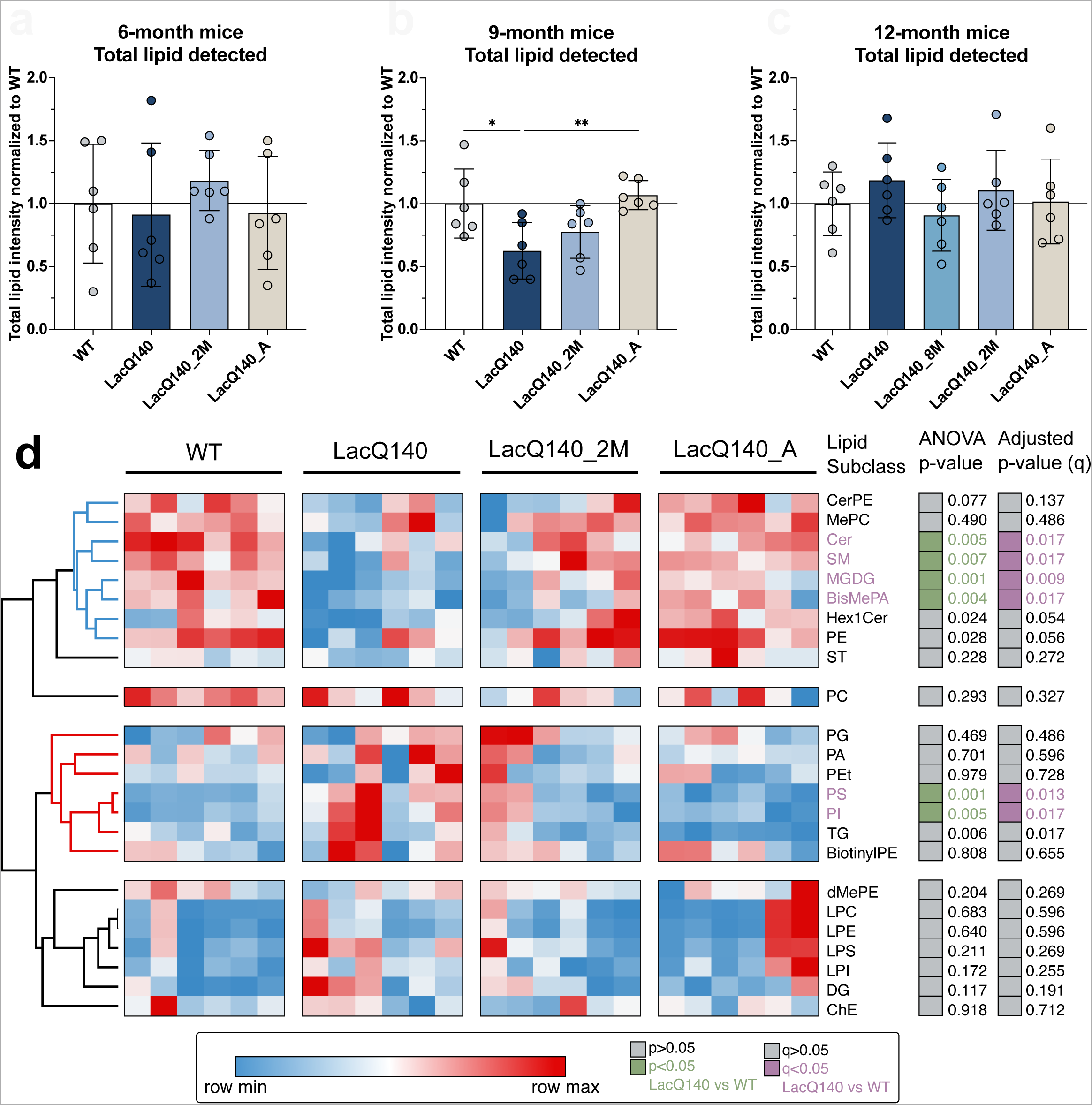
Analyses of lipids in crude homogenates LacQ140 caudate putamen by mass spectrometry. **(a)** Total lipid intensity detected at 6 months normalized to WT; no significant difference between groups (one-way ANOVA: F(3, 20) = 0.4604, P=0.7130, n.s., n=6). **(b)** Total lipid intensity detected at 9 months normalized to WT; LacQ140 mice have decreased total lipid intensity which is reversed in LacQ140_A mice (one-way ANOVA & Tukey’s multiple comparison test: F(3, 20) = 5.474, **P=0.0065, n=6). **(c)** Total lipid intensity detected at 12 months normalized to WT; no significant difference between groups (one-way ANOVA: F (4, 25) = 0.7504, P=0.5671, n.s., n=6). **(d)** Heatmap depicts the lipid subclass composition for WT, LacQ140, and treatment groups at 9 months. Hierarchical clustering was performed across lipid subclasses (rows) and columns (animals) using the one minus Pearson correlation distance metric. ANOVA p-value column indicates lipid subclasses significantly changed between LacQ140 and WT mice in green (p<0.05, One-way ANOVA with Tukey’s multiple comparison test, n=6). Lipid subclasses with adjusted p-values (q) < 0.05 LacQ140 vs WT are indicated in purple (q<0.05, two-stage linear step-up procedure of Benjamini, Krieger, and Yekutieli, N=24 lipid subclasses, n=6 mice). Source data and full statistical details can be found in **Additional Files 1, 4, & 5**.

**Table 1.** Overview of lipid subclass changes at 6, 9, and 12 months.

|  |  | 6-month-old mice |  |  | 9-month-old mice |  |  | 12-month-old mice |  |  |
| --- | --- | --- | --- | --- | --- | --- | --- | --- | --- | --- |
| Category | Subclass | Species detected | ANOVA p-val | Adj p-val (q), N =36 | Species detected | ANOVA p-val | Adj p-val (q), N =24 | Species detected | ANOVA p-val | Adj p-val (q), N =29 |
| Glycerophospholipids | PC | 134 | 0.7415 | 1 | 107 | 0.2928 | 0.3267 | 160 | 0.4243 | 0.4627 |
|  | PE | 152 | 0.1975 | 1 | 154 | 0.0284* | 0.0563 | 119 | 0.5653 | 0.5372 |
|  | PS | 62 | 0.1171 | 1 | 58 | **0.0014** | **0.0125** | 25 | 0.46 | 0.4627 |
|  | PI | 19 | **0.0189** | 0.7144 | 17 | **0.0054** | **0.0168** | 34 | 0.0086* | 0.0338* |
|  | MePC | 17 | 0.9859 | 1 | 7 | 0.4901 | 0.486 | 15 | 0.7552 | 0.6329 |
|  | PA | 13 | 0.9596 | 1 | 6 | 0.7008 | 0.5957 | 9 | 0.0025* | 0.0184* |
|  | BisMePA | 4 | 0.1589 | 1 | 1 | **0.0041** | **0.0168** | 6 | 0.2812 | 0.3647 |
|  | dMePE | 9 | 0.195 | 1 | 2 | 0.2041 | 0.269 | 0 | N/A | N/A |
|  | PG | 19 | 0.6255 | 1 | 4 | 0.4685 | 0.486 | 4 | 0.859 | 0.6765 |
|  | BisMePE | 1 | 0.8072 | 1 | 0 | N/A | N/A | 0 | N/A | N/A |
|  | BisMePS | 0 | N/A | N/A | 0 | N/A | N/A | 2 | 0.0001* | 0.0022* |
|  | CL | 22 | 0.5812 | 1 | 0 | N/A | N/A | 0 | N/A | N/A |
|  | PEt | 7 | 0.3261 | 1 | 2 | 0.9792 | 0.7283 | 0 | N/A | N/A |
|  | PMe | 2 | 0.8695 | 1 | 0 | N/A | N/A | 0 | N/A | N/A |
|  | PIP2 | 2 | 0.1393 | 1 | 0 | N/A | N/A | 3 | 0.0161* | 0.0394* |
|  | PIP | 1 | 0.7034 | 1 | 0 | N/A | N/A | 0 | N/A | N/A |
|  | LPC | 30 | 0.7943 | 1 | 10 | 0.6835 | 0.5957 | 7 | 0.2805 | 0.3647 |
|  | LPE | 23 | 0.9947 | 1 | 16 | 0.6396 | 0.5957 | 4 | 0.0127* | 0.035* |
|  | LPS | 7 | 0.9061 | 1 | 5 | 0.211 | 0.269 | 2 | 0.0817 | 0.1287 |
|  | LPI | 7 | 0.7746 | 1 | 1 | 0.1715 | 0.2551 | 0 | N/A | N/A |
|  | LPG | 4 | 0.8888 | 1 | 0 | N/A | N/A | 0 | N/A | N/A |
|  | LdMePE | 1 | 0.4552 | 1 | 0 | N/A | N/A | 0 | N/A | N/A |
|  | BiotinylPE | 0 | N/A | N/A | 1 | 0.8078 | 0.6554 | 1 | 0.0225* | 0.0451* |
| Glycerolipids | TG | 107 | 0.5998 | 1 | 44 | 0.0057* | 0.0168* | 72 | 0.0002* | 0.0022* |
|  | MGDG | 3 | 0.8693 | 1 | 2 | **0.0005** | **0.0089** | 5 | 0.0214* | 0.0451* |
|  | MGMG | 6 | 0.7537 | 1 | 0 | N/A | N/A | 6 | 0.0497* | 0.0887 |
|  | DG | 13 | 0.1381 | 1 | 70 | 0.1174 | 0.1905 | 59 | 0.6614 | 0.5834 |
|  | SQMG | 2 | 0.5986 | 1 | 0 | N/A | N/A | 0 | N/A | N/A |
|  | SQDG | 1 | 0.9316 | 1 | 0 | N/A | N/A | 0 | N/A | N/A |
| Sphingolipids | Hex1Cer | 0 | N/A | N/A | 85 | 0.0243* | 0.0542 | 104 | 0.4617 | 0.4627 |
|  | CerG1 | 50 | 0.5466 | 1 | 0 | N/A | N/A | 0 | N/A | N/A |
|  | SM | 47 | 0.6398 | 1 | 23 | **0.0066** | **0.0168** | 36 | 0.3243 | 0.3973 |
|  | Cer | 14 | 0.9804 | 1 | 6 | **0.0048** | **0.0168** | 21 | 0.4374 | 0.4627 |
|  | CerP | 0 | N/A | N/A | 0 | N/A | N/A | 17 | 0.0035* | 0.0193* |
|  | ST | 10 | 0.3977 | 1 | 9 | 0.2283 | 0.2717 | 14 | 0.2176 | 0.3199 |
|  | So | 3 | 0.4465 | 1 | 0 | N/A | N/A | 0 | N/A | N/A |
|  | CerPE | 0 | N/A | N/A | 1 | 0.0766 | 0.1367 | 0 | N/A | N/A |
|  | phSM | 4 | 0.3619 | 1 | 0 | N/A | N/A | 0 | N/A | N/A |
|  | CerG2GNAc1 | 1 | 0.5784 | 1 | 0 | N/A | N/A | 0 | N/A | N/A |
|  | SPH | 0 | N/A | N/A | 0 | N/A | N/A | 1 | 0.775 | 0.6329 |
|  | Hex2Cer | 0 | N/A | N/A | 0 | N/A | N/A | 1 | 0.0112 | 0.035 |
| Sterol Lipids | ChE | 1 | 0.5341 | 1 | 1 | 0.918 | 0.7124 | 2 | 0.9151 | 0.6958 |
|  | ZyE | 0 | N/A | N/A | 0 | N/A | N/A | 3 | 0.0092* | 0.0338* |
| Fatty Acyls | FA | 2 | 0.5893 | 1 | 0 | N/A | N/A | 0 | N/A | N/A |
|  | AcCa | 0 | N/A | N/A | 0 | N/A | N/A | 2 | 0.5847 | 0.5372 |
| Prenol Lipids | Co | 0 | N/A | N/A | 0 | N/A | N/A | 1 | 0.0523 | 0.0887 |
| Total |  | 800 |  |  | 632 |  |  | 735 |  |  |

**Table 2.** Overall comparison of lipidomic results across ages.

| Age | # Species detected | # Species different at $p < 0.05^*$ | # Species different for WT/LacQ140 at $p < 0.05$ | # Species different at $q < 0.05^{**}$ | # Species different for WT/LacQ140 at $q < 0.05$ | # Species recovered with lowering ($q < 0.05$) |
| --- | --- | --- | --- | --- | --- | --- |
| 6 months | 800 | 9 | 4 | 0 | 0 | 0 |
| 9 months | 632 | 192 | 72 | 17 | 14 | 14 |
| 12 months | 735 | 162 | 36 | 72 | 26 | 1 |

\*p value for ANOVA; \*\*Adjusted p value determined using Benjamini, Krieger, and Yekutieli procedure

We assessed if there were more subtle alterations at the subclass or individual species level that did not influence the overall lipid levels. In striatum of LacQ140 mice at 6-months-of-age, we detected a modest increase in the subclass phosphatidylinositol (PI) compared to WT (**Fig. S7**). Individual species of PI (18:0_20:4, 16:0_22:6), and phosphatidylserine PS (18:0_20:4, 22:6_22:6) were increased in LacQ140 mice compared to WT (**Fig. S8**). However, no subclass or species changes survived correction using the Benjamini, Krieger, and Yekutieli procedure with a 5% false discovery rate (FDR) of q <0.05, N=36 subclasses (**Table 1**) and N=800 species (**Additional File 1)**. Consistent with our observations at 6 months, PS and PI were increased in 9-month-old LacQ140 mice compared to WT (**Fig. 5a and b**). A limited number of species are modestly increased at 6 months, which progresses to more robust increases in these glycerophospholipids at the subclass level at 9 months. A significant reduction in bismethyl phosphatidic acid (BisMePA) also occurred (**Fig. 5c**). Functionally, PI is the precursor for PIPs which are important for protein kinase c (PKC) signaling at the synapse ^85^ and can also act as important docking and activating molecules for membrane associated proteins, including HTT ^52^. PS is an abundant anionic glycerophospholipid necessary for activation of several ion channels ^86^, fusion of neurotransmitter vesicles, regulation of AMPA signaling, and coordination of PKC, Raf-1, and AKT signaling ^87^.

**Figure 5.**
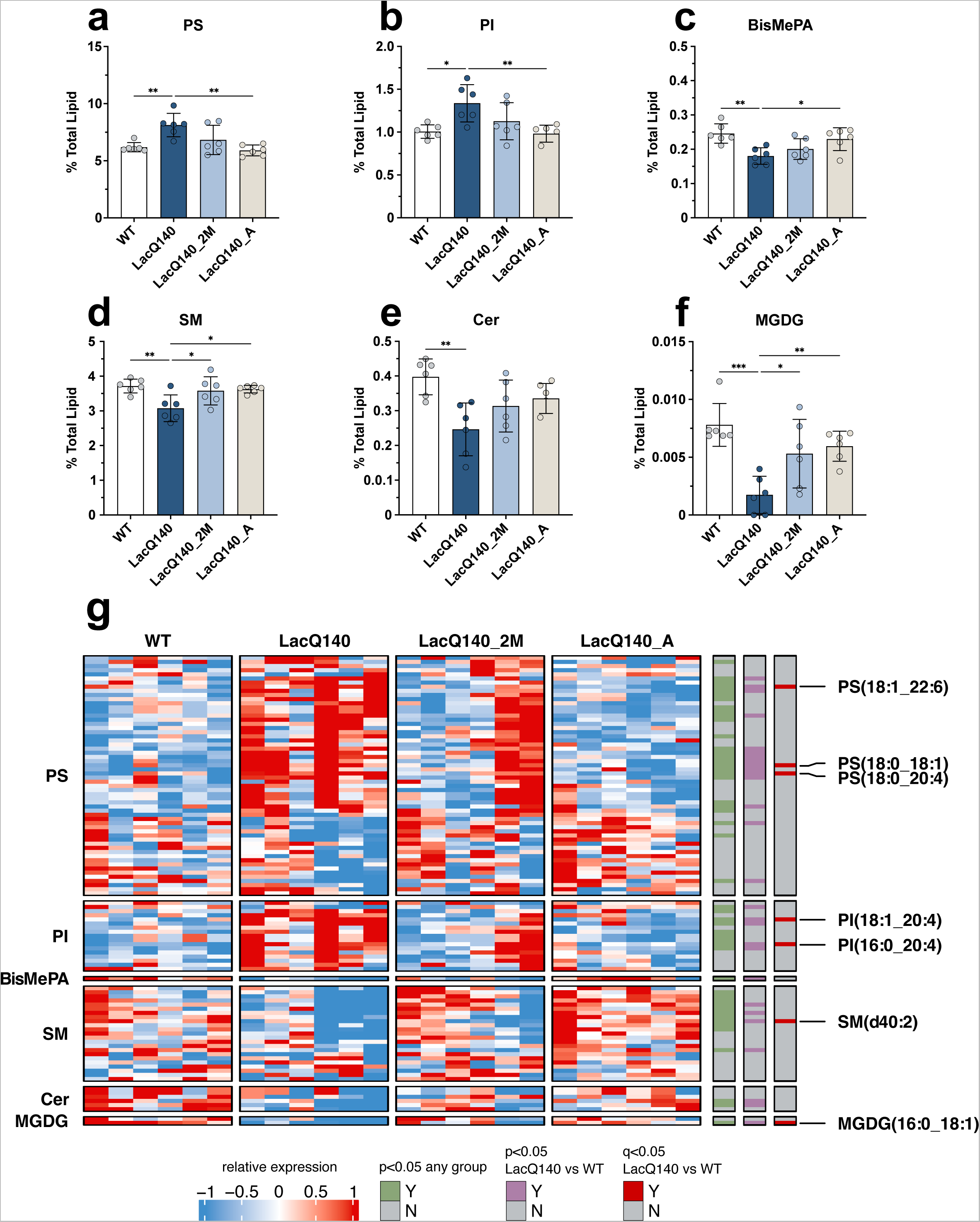
Restoration of dysregulated lipid subclasses with lowering of m*Htt*. Graphs show relative intensities for lipid subclasses expressed as a percent of total lipid intensity detected (bars = mean, error bars = ± SD). **(a)** PS increased in LacQ140 mice and is reversed in LacQ140_A mice; F(3, 20) = 7.601, **P=0.0014, q=0.0125, **(b)** PI increased in LacQ140 mice and reversed in LacQ140_A mice; F(3, 20) = 5.707, **P=0.0054, q=0.0168, **(c)** BisMePA decreased in LacQ140 mice and is reversed in LacQ140 mice; F(3, 20) = 6.086, **P=0.0041, q=0.0168, **(d)** SM decreased in LacQ140 mice and is reversed in LacQ140_2M and LacQ140_A mice; F(3, 20) = 5.465, **P=0.0066, q=0.0168, **(e)** Cer decreased in LacQ140 mice; F(3, 20) = 5.883, **P=0.0048, q=0.0168, **(f)** MGDG decreased in LacQ140 mice and is reversed in LacQ140_2M and LacQ140_A mice; F(3, 20) = 9.350, ***P=0.0005, q=0.0089. Statistics are one-way ANOVA and asterisks on graphs represent Tukey’s multiple comparison test, n=6 mice per group. **(g)** Heatmap shows individual lipid species that comprise each subclass. Hierarchical clustering was performed across individual lipid species (rows) and animals (columns) using the one minus Pearson correlation distance metric. Individual lipid species significantly changed between any group are indicated in green (p<0.05, one-way ANOVA), lipid species significantly changed between LacQ140 and WT are indicated in purple (p<0.05, one-way ANOVA, n=6) and red (q<0.05, one-way ANOVA & two-stage linear step-up procedure of Benjamini, Krieger, and Yekutieli, N=632 lipid species, n=6 mice). Source data and full statistical details can be found in **Additional Files 1, 4, & 5**.

In contrast to the increases observed in glycerophospholipids, 9-month-old LacQ140 mice exhibit significant reductions in the subclasses sphingomyelin (SM) and ceramide (Cer) (**Fig. 5d and e**), lipids central in sphingolipid metabolism and important for myelination. Cer is the backbone for SM as well as more complex glycosphingolipids, which are highly enriched in myelin ^88^. Moreover, the low abundance glycerolipid monogalactosyldiacylglycerol (MGDG) was strikingly reduced at 9 months (**Figure 5f**). MGDG regulates oligodendrocyte differentiation and is considered a marker of myelination ^89,90^.

All subclasses (PI, PS, BisMePA, SM, Cer, MGDG) with changes between WT and LacQ140 had q values<0.05 (N=24 subclasses) (**Table 1**; **Fig. 4d**). Early lowering of m*Htt* (LacQ140_A) restored levels of each of these subclasses except for Cer and lowering m*Htt* starting at 2 months (LacQ140_2M) was sufficient to correct levels of SM and MGDG (**Fig. 5a-f**). To assess the effects of m*Htt* lowering on the distinct lipid species that comprise each subclass, lipid species were plotted in a heatmap (**Fig. 5g**). The broad changes at the subclass level can be attributed to large numbers of individual lipid species changing within each subclass even if each species is not significantly different (**Fig. 5g** & **Fig. S9**). Alternatively, subclasses can remain unchanged but still contain individual lipid species with differences. For example, the subclass Hex1Cer was not significantly changed **(Fig. S9)**, but many individual species were significantly altered in LacQ140 mice (**Fig. S10**). These species included 3 that were significantly decreased in LacQ140 mice and normalized to WT levels with lowering of m*Htt*: Hex1Cer (d18:1_18:0), Hex1Cer (t18:0_18:0), Hex1Cer (d18:1_25:1) (**Fig. S10**). Combined, these 3 species represent approximately 1% of total Hex1Cer detected. Hex1Cer is comprised of galactosylceramide and glucosylceramide and cannot be distinguished by LC-MS/MS; however, previous studies have determined that the majority of Hex1Cer in the brain is galactosylceramide, which is highly enriched in myelin ^67,91^. Galactosylceramides are major constituents of the myelin sheath and contribute to its stabilization and maintenance ^92^.

In total, we identified 14 lipid species significantly altered between LacQ140 and WT mice using a stringent FDR cutoff (FDR 5%/q<0.05, Benjamini, Krieger, and Yekutieli procedure, N=632 lipids) and all 14 showed improvement with m*Htt* lowering at 9 months (**Fig. S10**). This included increases in PI (16:0_20:4) & PI (18:1_20:4) and PS (18:0_20:4), PS (18:0_18:1) and PS (18:1_22:6). Lipid species decreased in LacQ140 mice and restored with m*Htt* lowering included Hex1Cer species as detailed above, SM (d40:2), PE (16:0p_20:4) and MGDG (16:0_18:1) (**Fig. 5g**, **Fig. S10**). Taken together, these results indicate a profound disruption of myelin lipids in LacQ140 mice, which is attenuated with lowering of m*Htt*.

In the striatum of LacQ140 mice at 12 months, no significant differences at the subclass level were found compared to WT (**Table 1**; **Fig. S11**) although 29 subclasses were measured. At the individual lipid species level, we identified 26 lipid species significantly altered between LacQ140 and WT mice (FDR 5%/q<0.05, Benjamini, Krieger, and Yekutieli procedure, N=735 lipids). (**Fig. S12**). Contrary to the reversal of lipid dysregulation observed at 9 months, the only lipid species that was responsive to lowering m*Htt* at 12 months was CerP (t40:3), which was increased in LacQ140 and decreased with early m*Htt* lowering (LacQ140_A) (**Fig. S12**). Myelin enriched lipids, including 6 species of Hex1Cer (d18:1_18:0, d:40:2, d18:2_22:0, d41:2, t18:0_22:1 & d18:1_22:1) and ST (d18:1_22:1), decreased and were not improved with m*Htt* lowering (**Fig. S12**). Similarly, nine triacylglycerol (TG) species, eight of which contain oleic acid (C18:1) were decreased in LacQ140 mice and unchanged with m*Htt* lowering (**Fig. S12**). Overall, these results demonstrate that most individual lipid changes were no longer reversible by 12 months.

### Bioinformatic analysis of transcriptional alterations to lipid metabolism and myelination

To determine if any of the lipid changes could be explained by altered transcription of genes regulating lipid-related metabolic pathways, we re-analyzed the previously generated RNAseq dataset in the LacQ140 striatum (GEO GSE156236). Differential expression analysis ^70^ detected 1360 upregulated and 1431 downregulated genes at 6 months and 1375 upregulated and 1654 downregulated genes at 12 months (FC>20%, adjusted p-value <0.05) (**Fig. S13a**). Over-representation analysis against GO Biological Processes (GO BP) revealed that downregulated genes were enriched in terms related to lipid metabolism and signaling as well as myelination. In 6-month LacQ140 downregulated genes, GO BP terms ensheathment of neurons (padj<0.01), axon ensheathment (padj<0.01), myelination (padj<0.05), cellular lipid metabolic process (padj<0.05), lipid metabolic process (padj<0.05), phospholipase C-activating G protein-coupled receptor signaling pathway (padj<0.05), response to lipid (padj<0.05) were significantly over-represented (**Fig. 6a**). In LacQ140 upregulated genes, there was no over-representation of terms that could directly explain changes to lipid levels **(Additional File 2).**

**Figure 6.**
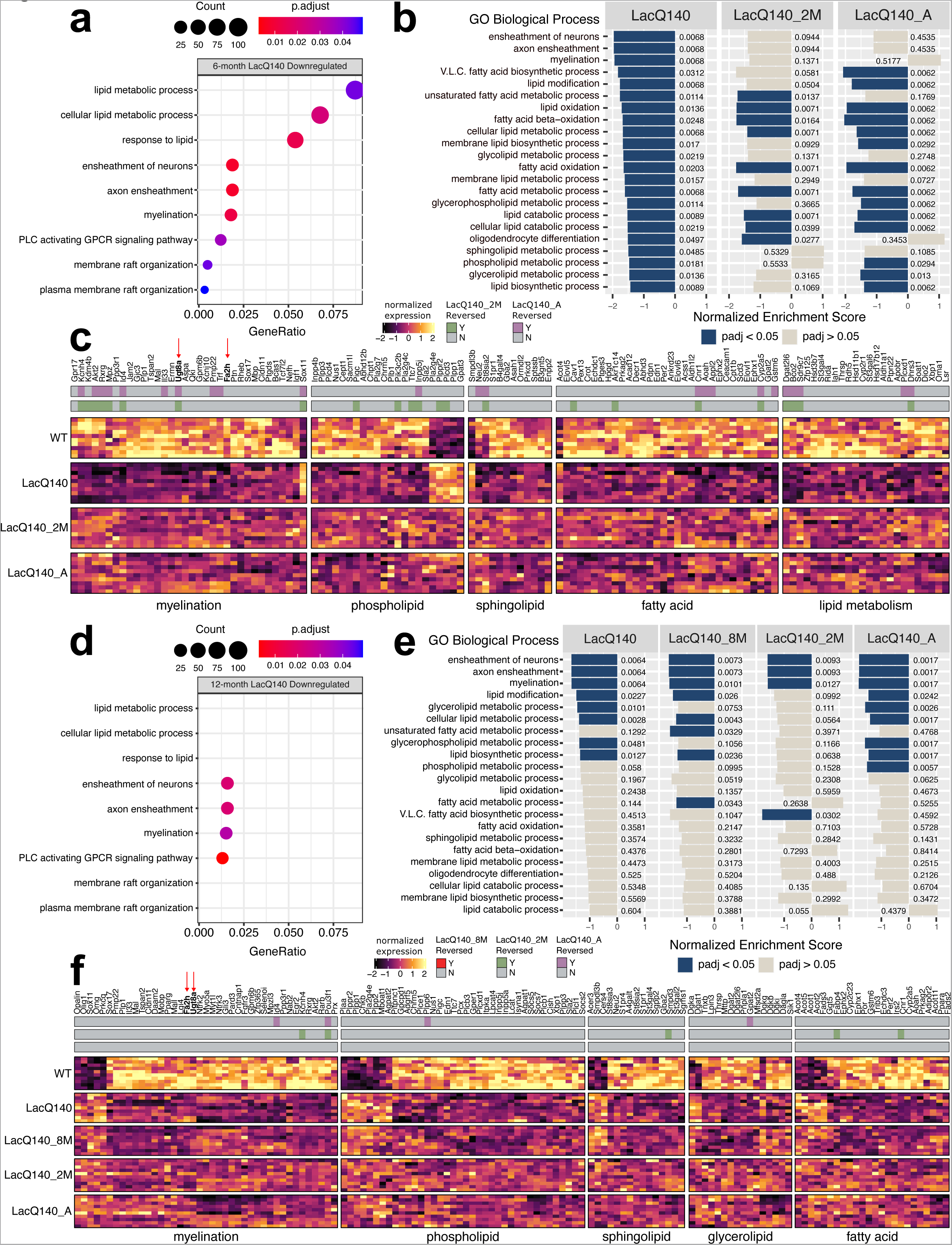
Lipid metabolism and myelin associated transcriptional changes and reversal with m*Htt* lowering. **(a)** Dotplot (clusterProfiler) of lipid or myelin related GO BP terms overrepresented (one sided hypergeometric test, padj<0.05) in 6-month LacQ140 downregulated genes (padj<0.05, FC>20%). GeneRatio represents the number of genes associated with each GO term/number of downregulated genes. Dots are sized by count of genes associated with respective terms. **(b)** Gene set enrichment analysis (clusterProfiler) of 6-month LacQ140, LacQ140_2M and LacQ140_A groups compared to WT. All lipid and myelin associated GO BP terms significantly enriched in LacQ140 compared to WT (padj<0.05) are displayed. X axis for each respective group represents the normalized enrichment score (NES) and bars are colored by significance (padj<0.05 = blue, padj>0.05 = tan). Adjusted p-values are displayed adjacent to bars. **(c)** Heatmap shows differentially expressed genes in 6-month-old LacQ140 mice compared to WT (padj<0.05, FC > ±20%). Gene expression is shown as median ratio normalized counts (DESeq2), scaled by respective gene (columns). Bars above the heatmap indicate DEGs reversed with m*Htt* lowering. LacQ140_2M = green, LacQ140_A = purple; padj<0.05, FC > 20% opposite of LacQ140**. (d)** Dotplot of lipid or myelin related GO BP terms overrepresented in 12-month downregulated genes. GO BP terms significantly overrepresented at 6-months (a) are displayed for comparison. 4/9 terms (ensheathment of neurons, axon ensheathment, myelination, and phospholipase C activating G-protein coupled receptor signaling pathway) are significantly overrepresented in 12- month LacQ140 downregulated genes (one sided hypergeometric test, padj < 0.05). **(e)** Gene set enrichment analysis of LacQ140, LacQ140_8M, LacQ140_2M and LacQ140_A groups compared to WT. GO BP terms enriched at 6-months are displayed for comparison. 8/22 GO BP terms are significantly negatively enriched (padj < 0.05) at 12-months, shown in blue. **(f)** Heatmap shows differentially expressed genes in 12-month-old LacQ140 mice compared to WT (padj<0.05, FC > ±20%). Gene expression is shown as median ratio normalized counts (DESeq2), scaled by respective gene (columns). Bars above the heatmap DEGs reversed with m*Htt* lowering (LacQ140_8M = bottom bar, no reversal, LacQ140_2M = green, LacQ140_A = purple; padj<0.05, FC >20% opposite of LacQ140). GO BP terms associated with genes, fold changes, and exact FDR values can be found in **Additional Files 2 & 3.**

Next, Gene Set Enrichment Analysis (GSEA) was conducted ^93,94^ to examine the impact of m*Htt* repression on the enrichment profiles of terms related to lipid metabolism and myelination. At 6 months, 3 of the top 20 enriched GO BP terms were related to myelination and all were negatively enriched: ensheathment of neurons, myelination, and axon ensheathment **(Additional File 2)**. Further, every significantly enriched myelin or lipid related GO BP term was negatively enriched in LacQ140 compared to WT (padj <0.05) (**Fig. 6b**, **Additional File 2)**. Repression of m*Htt* in LacQ140_2M and LacQ140_A groups was sufficient to reverse the negative enrichment of many GO BP terms including myelination/axon ensheathment/ensheathment of neurons, and sphingolipid/phospholipid metabolic processes suggesting some beneficial impact of m*Htt* lowering (**Fig. 6b**).

Specifically, 6-month LacQ140 mice had reduced expression of genes encoding proteins involved in oligodendrocyte development (*Akt2, Bcas1, Ptgds, Cldn11, Trf, Tcf7l2, Qki*), myelin structure and compaction (*Mbp, Plp1, Tspan2, Mal*), and myelin lipid biosynthesis (*Ugt8a, Fa2h, Aspa*) (**Fig. 6c**). LacQ140 mice also exhibited dysregulation of genes encoding enzymes including phospholipases (*Pla2g7, Pla2g4c, Pla2g4e, Plb1, Plcd3, Plcd4*), phosphatases & kinases (*Plpp1, Plpp3, Plppr2, Inpp4b, Pik3c2b*), and regulators of phospholipid biosynthesis (*Gpat2, Gpat3, Chpt1*) (**Fig. 6c**). Enzymes responsible for biosynthesis and metabolism of sphingolipids were altered in

LacQ140 mice: *Sptssb, Ormdl2, Gba2, Asah,* were downregulated and *Smpdl3b* was upregulated. Genes encoding enzymes responsible for fatty acid elongation (*Elovl1, Elovl5, Elovl6*) were all downregulated (**Fig. 6c**). Of the DEGs related to lipid metabolism and myelination 23 were reversed in LacQ140_2M mice, 23 were reversed in LacQ140_A mice and 15 were reversed in both (padj < 0.05, FC > 20% opposite direction of LacQ140) (**Fig. 6c** & **Fig. S13b**).

Three of the nine GO BP terms significantly over-represented in 6-month LacQ140 downregulated genes were also over-represented in 12-month downregulated genes: ensheathment of neurons (padj <0.01), axon ensheathment (padj <0.01), myelination (padj <0.05), and phospholipase C-activating G protein-coupled receptor signaling pathway (padj<0.01) (**Fig. 6d**). Similarly, GSEA showed that only a subset (8/22) of the GO BP terms related to lipid metabolism or myelination were enriched at both timepoints (**Fig. 6e**). Terms including glycerophospholipid metabolic process and glycerolipid metabolic processes showed improvement with lowering of m*Htt*, whereas terms related to myelination remained negatively enriched with m*Htt* repression (**Fig. 6e**).

DEGs associated with myelin and lipid biosynthesis were both downregulated and upregulated in 12-month LacQ140 mice. LacQ140 mice had reduced (*Cntnap1, Mal, Tspan2, Plp1, Mobp*) and increased (*Omg, Opalin*) expression of genes encoding proteins involved in myelin structure and compaction (**Fig. 6f**). Similarly, oligodendrocyte development related genes were both decreased (*Akt2, Cldn11, Myt1l*) and increased (*Prkcq, Olig1, Sox11*). Genes related to myelin lipid biosynthesis (down: *Ugt8a, Fa2h, Selenoi*), phospholipid synthesis/metabolism (up: *Chkb, Agpat2, Mboat2,* down: *Agpat1, Lpcat4*), phosphatases (up: *Plpp1, Plpp2, Plppr2,* down: *Plpp3, Plpp6, Inpp5a*), phospholipases (up: *Pla2g4e,* down: *Plcb1, Plcb3, Plce1, Plcl1*) and sphingolipid synthesis/metabolism (up: *Cers4, Smpdl3b, Acer3,* down: *Gba2, St3gal2, Smpd3, Sgpp2, Sgms1, A4galt, Fads3*) were differentially expressed in both directions (**Fig. 6f**). In 12-month LacQ140 mice, glycerolipid synthesis/metabolism genes (up: *Dgat1,* down: *Dgat2, Dgat2l6, Lpin3, Dagla*) and DAG kinase genes (up: *Dgka, Dgkg,* down: *Dgkb, Dgki*) were differentially expressed (**Fig. 6f**). Consistent with the enrichment results at this timepoint, fewer genes were reversed with repression of m*Htt* (5 genes LacQ140_2M, 4 genes LacQ140_A) (**Fig. 6f** & **Fig. S13c**).

Differentially expressed genes (DEGs) were also manually curated for genes that could directly impact levels of PS, PI, sphingolipids, and glycerolipids. Phosphatidylserine synthase enzymes (*Ptdss1/Ptdss2*) ^87^ and rate limiting enzymes for PI biosynthesis (CDP-Diacylglycerol Synthases: *Cds1/Cds2*) ^95^ were unchanged in 6- and 12-month LacQ140 mice **(Additional Files 1 & 2)**. For glycerolipids, the only known enzymes that catalyze TG biosynthesis ^96^ were increased (*Dgat1*) and decreased (*Dgat2*) at 12 months; mRNA levels remained unchanged with m*Htt* repression (**Fig. 6f**).

Lipid changes might be explained by altered cellular composition of tissue. To address this possibility, we surveyed for changes in transcript levels for cellular markers. No change in mRNA expression levels for the microglial markers *Iba1* and *Cd68* or for the reactive astrocyte marker *Gfap* were observed that might indicate upregulation of these cell types could account for the lipid changes. However, our transcriptional profiling did reveal an expression signature consistent with altered oligodendrocyte development and alterations to myelin structure or compaction in both 6- and 12-month-old mice (**Fig. 6c and f**). UDP-galactose-ceramide galactosyltransferase (*Ugt8a*) is expressed in oligodendrocytes where it catalyzes the formation of GalCer (Hex1Cer) from Cer; *Ugt8a* mRNA was decreased at both 6 and 12 months in the LacQ140 mice (**Fig. 6c** & **Fig. 7a**). Embryonic repression of m*Htt* and repression beginning at 2-months of age preserved *Ugt8a* mRNA measured at 6-months; however, the effect was lost by 12-months of age (**Fig. 6c and f**). These mRNA results mirror our findings of Hex1Cer lipid levels measured using LC-MS in LacQ140 striatal lysates. GalCer and GlcCer are both members of the Hex1Cer subclass but cannot be distinguished by MS since they are isomers with the same mass and charge ^67^; the gene catalyzing the transfer of glucose to ceramide (*Ugcg*) was not changed at the mRNA level, suggesting lower levels of Hex1Cer in LacQ140 mice are due to lower levels of GalCer. Ugt8a can also transfer galactose to diacylglycerol to form the lipid MGDG ^97^ which we found was decreased with m*Htt* expression and improved with m*Htt* lowering (**Fig. 5f and g**). In contrast, *Fa2h* mRNA was reduced at 6 and 12-months but not improved with *mHtt* lowering (**Fig. 6c and f**). Fatty acid 2-hydroxylase, which is encoded by *Fa2h,* modifies fatty acids substrates with a hydroxyl (-OH) group, producing 2-hydroxylated fatty acids that are incorporated into myelin sphingolipids and provide stability^98^ (**Fig. 7b and c**). Overall, these data reveal transcriptomic changes in key lipid metabolic and biosynthetic genes in LacQ140 striatum, which directly impact the lipidomic profile. Critically, many of these lipidomic and transcriptomic alterations can be attenuated with lowering of m*Htt*.

**Figure 7.**
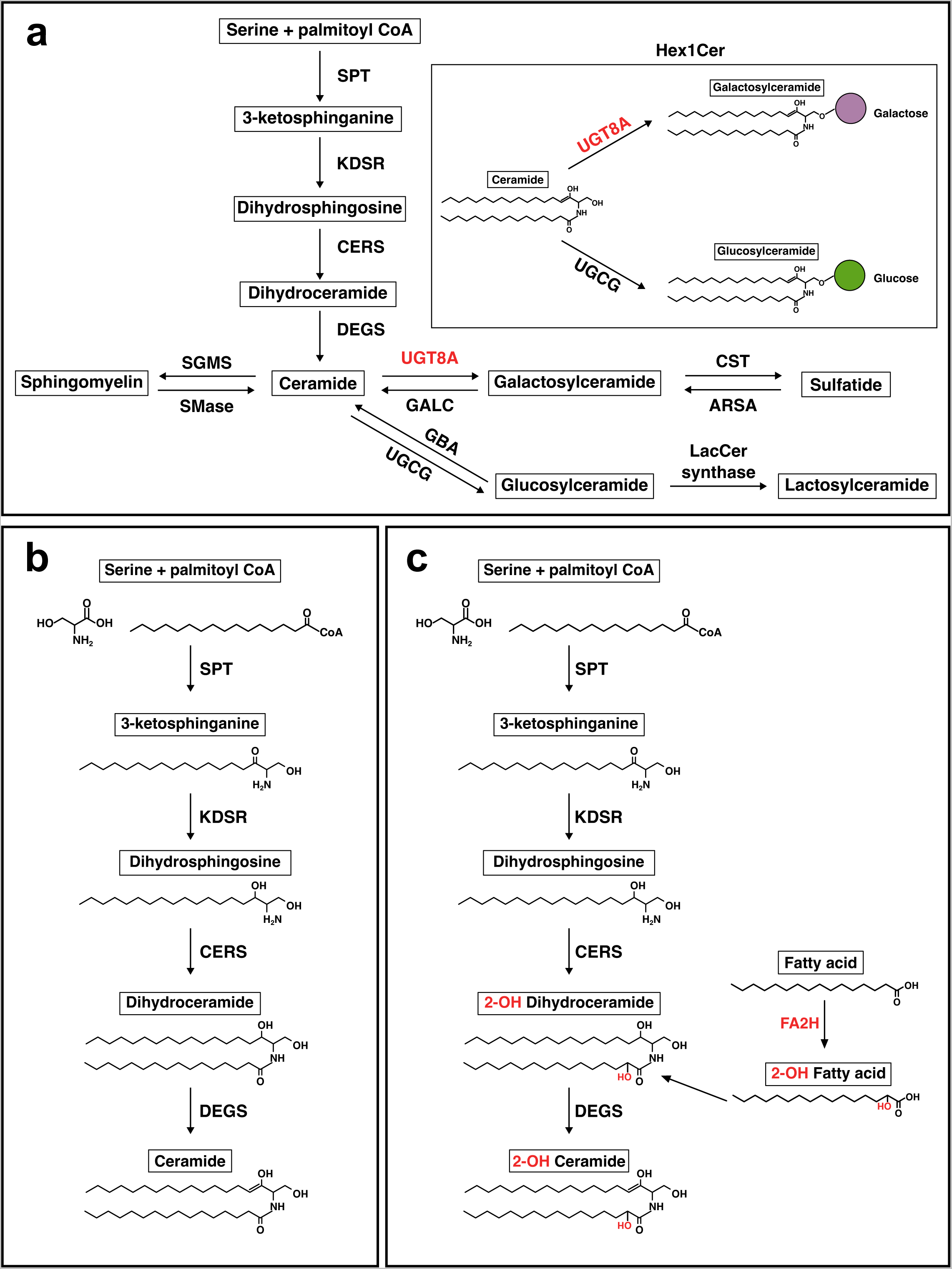
Simplified de novo sphingolipid biosynthesis pathway. **(a)** Serine and palmitoyl CoA are condensed by serine palmitoyltransferase (SPT) to generate 3-ketosphinganine. 3- ketophinganine is reduced to dihydrosphingosine by 3-ketodihydrosphingosine reductase (KDSR). Dihydrosphingosine is acetylated by ceramide synthases (CERS) and further desaturated by ceramide desaturase (DEGS) to generate ceramide. Ceramide is the substrate for generation of other sphingolipids (sphingomyelin, galactosylceramide, glucosylceramide, sulfatide, and lactosylceramide). Abbreviations: SGMS = sphingomyelin synthase, SMase = sphingomyelinase, UGT8A = UDP galactosyltransferase 8A, GALC = galactosylceramidase. CST = galactosylceramide sulfotransferase, ARSA = arylsulfatase, GBA = glucosylceramidase, UGCG = UDP-glucose ceramide glucosyltransferase, LacCer synthase = lactosylceramide synthase. **(b)** Biosynthesis of non-hydroxylated sphingolipids. Biosynthesis of 2-hydroxy sphingolipids. Fatty acid 2-hydroxylase (FA2H) catalyzes hydroxylation of fatty acids in the C2 position, which can be incorporated into sphingolipid precursors (i.e., dihydroceramide) in the acylation step of de novo synthesis.

## DISCUSSION

Here we used the LacQ140 inducible HD mouse model to initiate whole body m*Htt* reduction at different ages and evaluate effects on proteins and lipids. Lowering mHTT protein by 38-52% in the LacQ140 caudate putamen, starting from conception up to 12 months of age, was sufficient to prevent m*Htt* induced changes in the levels of some proteins and some lipids. However, a resistant soluble species of the protein detected in older mice limited long term benefit of m*Htt* lowering. Our lipid data show clear evidence of changes impacting myelin which mechanistically is due in part to aberrant transcription as evidenced by differentially expressed genes regulating some of the same lipids that were found altered. The correction of lipid changes we identified here correlates with the behavioral changes measured previously in LacQ140 mice using a comprehensive set of unbiased, high-content assays ^59^ in that behavior was closer to normal at 6 and 9 months with early mHTT lowering, but the protective effect of lowering degraded by 12 months.

We identified forms of mHTT protein that were resistant to m*Htt* lowering detected by various HTT antibodies. These resistant forms of mHTT identified by immunoblot may correspond to mHTT aggregates or foci found to be resistant to lowering in LacQ140 mice using MSD and immunofluorescence methods ^59^. Others have described aggregated and soluble forms of mHTT that resist degradation in distinct cellular compartments, including full length mHTT in the nucleus ^17,99–101^. In the LacQ140 striatum, the SDS-soluble degradation-resistant form of full-length mHTT, detected by us using antibodies S830, MW1 and EPR5526, resides in a perinuclear or nuclear compartment which might correspond to the “juxtanuclear quality control compartment (JUNQ)” described by Frydman and colleagues ^102^. Accrual of misfolded mHTT in the nucleus likely contributes to transcriptional interference and the eventual failure of prolonged benefits of modest mHTT lowering. Targeting these resistant fractions of misfolded mHTT by a chaperone activity to aid in its degradation may be beneficial in combination with gene therapy *HTT* lowering strategies.

We found that early, continuous partial lowering of mHTT protein for up to 12 months fully or partially preserved PDE10A and SCN4B protein levels with initiation of lowering embryonically providing the highest benefits. These data agree with previous findings showing preservation of *Pde10a* mRNA in the LacQ140 model after early m*Htt* lowering ^59^ and preservation of the Pde10a PET signal in the Q175 model after striatal injection of AAV-HTT-ZFP17. However, our data here indicate that the time points chosen for post-treatment analysis are important and that changes do not follow a linear neurodegenerative trajectory in mice. The greatest number of protein and lipid changes in LacQ140 mice occurred at 9 months, with few changes detected at 12 months. Similarly, Langfelder et al. found a greater number of differentially expressed genes at 6 months compared to 10 months in zQ175 mice ^39^. This suggests that in the mouse brain, adverse responses to mHTT oscillate or go through waves of degeneration and regeneration. Therefore, to appreciate any benefits afforded by m*Htt* lowering, frequent or continuous monitoring should be conducted.

In this study, mass spectrometry of lipids identified numerous alterations in LacQ140 striatum, many of which were prevented with modest m*Htt* lowering. To our surprise, lipidomic analysis showed that LacQ140 mice had increased levels in species of glycerophospholipids PI and PS starting at 6 months and progressing to a significant change in the PI and PS subclass level at 9 months. Proteomics in Q140 synaptosomes revealed changes in proteins that regulate PI levels including PKC signaling and PIP2 hydrolysis, changes in two isoforms of DGKs, and alterations in one of the rate-limiting enzymes in PI synthesis (CDS2) ^33^ all of which impact PI levels ^95^. Transcriptomic profiling of LacQ140 mice also showed a plethora of altered mRNA levels for enzymes that impact PI and PIPs (**Fig. 6c and f**; **Additional Files 2 and 3**). Normally found on the inner leaflet of the plasma membrane, PS can be externalized by apoptotic cells to signal for their demolition ^103^ and in neuronal synapses, externalized PS signals to microglia for synaptic pruning ^104^.Microglial activation occurs in HD post-mortem brain ^105^ and increased pruning by microglia is hypothesized to contribute to synaptic loss in R6/2 HD striatum ^106^ and in zQ175 and BACHD mice, as well as in HD brain ^107^. An overall increase in PS could inadvertently mark synapses or myelin ^108^ for engulfment by microglia. The ratio of PS to PE impacts autophagy ^109^ which may in turn impair mHTT removal ^110,111^. Both PI and PS are abundant in astrocytes as well as neurons ^112^ so it is unclear which cell type(s) is producing the changes in these lipids. We cannot rule out the possibility that lipid changes are due to altered cellular composition of the brain.

Changes in white matter detected through imaging are one of the first signs of disease in people with HD (PwHD) ^113–127^. Morphometric studies of postmortem HD brains showed reduced cross-sectional area of white matter as well as gray matter atrophy ^128,129^. In HD post-mortem brain tissue, a dramatic shift in the profile of various sphingolipids including Cer, SM, hexosylceramides, and sulfatides occurred ^51^. Here, we demonstrate in the striatum of 9-month-old LacQ140 mice significant reductions (compared to WT mice) of relative levels of total lipids and the lipid subclasses SM and Cer, and three individual species of Hex1Cer, all important for myelin ^130^. Our data are in alignment with data from R6/1 mice showing changes in cerebroside and sulfatide levels ^131^, from the R6/2 mouse model showing reductions in components of the sphingolipid biosynthesis pathway ^45^, and findings in a transgenic sheep model OVT73, similarly showing decreased levels of numerous species of SM ^132^. A salient finding was a profound reduction in LacQ140 striatum of the low abundance signaling lipid MGDG, which regulates oligodendrocyte differentiation ^89^. MGDG is considered a marker of myelination and stimulates PKC-alpha activity in oligodendrocytes to support process formation ^90^. We previously reported reduced levels of MGDG in subcellular fractions of Q175/Q7 HD striatum at 2 and 6 months ^47^. Crucially, in LacQ140 mice, lowering m*Htt* improved the loss of SM and MGDG suggesting protection against white matter pathology.

Altered oligodendrocyte differentiation or survival due to direct effects of mHTT protein on transcription in the nucleus may account for many of the lipid changes we observed. RNA transcripts of genes important for oligodendrocyte differentiation and myelin sphingolipid biosynthesis were altered in LacQ140 mice. The transcription factor *Tcf7l2*, which was lower at 6 months in LacQ140 mice, was recently implicated in altered myelin formation in R6/2 and Q175 mice ^133^. The basic helix-loop-helix transcription factor *Olig1*, which was increased at 12 months in LacQ140 mice compared to WT, is important for commitment of cells to CNS oligodendrocyte identity ^134^. Critically, Lim et al. presented evidence showing abnormal oligodendrocyte maturation in multiple HD postmortem brain regions, as well as R6/2 brain, with single-nucleus RNAseq showing changes in *OLIG1* and *OLIG2* ^135^. Altered levels of myelin transcripts were found in human embryonic stem cells differentiated along an oligodendrocyte pathway ^136^ and an epigenic etiology for changes in myelin gene expression in human oligodendrocyte precursors that was blocked by inactivation of m*HTT* allele was described ^137^.

Changes in particular enzyme levels that regulate lipid biosynthesis that were changed at the transcriptional level in LacQ140 mice can have dire consequences and result directly in myelin defects. Both *Ugt8a* and *Fa2h* mRNA were lower in the striatum of LacQ140 mice; *Ugt8a* mRNA levels were protected by m*Htt* lowering at 6 months, but *Fa2h* mRNA was not. Work by others showed that mice deficient in the *Ugt8a* gene exhibited abnormal myelin maturation and structure ^138,139^. In humans, mutations in FA2H are associated with leukodystrophy and hereditary spastic paraplegia type SPG35 ^140,141^ highlighting the importance of hydroxylated sphingolipids in myelin integrity. Adult *Fa2h-*deficient mice have normal oligodendrocyte differentiation with normal appearing myelin that later degenerates, showing “splitting of lammelae” by 18 months ^142^. This is similar to the Ki140CAG mouse model ^143^ where myelin appears to be quite normal into early adulthood, but then may start to degenerate with disease progression.

Although transcriptional deregulation clearly impacts lipid modifying enzymes, levels of myelin-related lipids could be caused by an interaction of mHTT with oligodendrocyte membranes or a be a consequence of Wallerian degeneration of cortical-striatal axons of the neurons. Interestingly, the presence of SM increased permeabilization of monolayers by mHTT *in vitro* ^53^, suggesting mHTT could have particular effects on myelin lipids. Crucially, mHTT was localized within myelin sheaths using immunogold EM in 9-month-old Q175 striatum ^80^. Moreover, mHTT can be secreted by neurons in culture and in the brain as a soluble free form ^144^. We speculate that mHTT could insert directly into myelin bilayers to disrupt myelin architecture.

Observing white matter changes in animal models has been challenging. White matter loss was reported in R6/1 mice ^145^, and changes in myelination have been described in Yac128 ^146,147^ and HdhQ250 mice ^148^, but experiments designed to look for white matter changes in the Q150 HD mouse model showed brain atrophy but no white matter abnormalities ^149^. A recent imaging study of OVT73 sheep brain reported changes in diffusivity in the internal capsule at 9-10 years, suggesting changes in white matter microstructure ^150^. Our biochemical experiments here show that mHTT effects on striatal lipid homeostasis in HD mouse models are complex. We and others have reported lipidomic and metabolomic studies on knock-in Q140/Q140 HD mice at single time points ^48^ and Q111 HD mice ^44^ but did not observe loss of lipids important for white matter. Curiously, the lipid differences in LacQ140 mice measured at 9 months disappeared at 12 months suggesting that, even in the absence of m*Htt* lowering, the mouse brain insulted with mHTT attempts to heal itself and succeeds at some level. Consistent with our lipidomic findings, a longitudinal imaging study over 18 months showed transient changes in diffusivity/fractional anisotropy of corpus collosum in Q140 mouse brain ^143^. These results echo imaging data from presymptomatic PwHD suggesting attempted remyelination ^151^. Hence, if HD mouse models undergo a series of degeneration and regeneration cycles, observations at one or two time points may be misleading. By 12 months, although changes at the lipid subclass level were annulled in the LacQ140 mice, a detailed analysis of the individual lipid species comprising these subclasses shows a shift in species within each subclass. The altered composition of subclasses may alter function, weaken HD brains, or predispose them to further stress. In fact, the behavioral analyses of these mice which showed loss of protection by mHTT lowering at 12 months ^59^ indicates that although levels of lipids by subclasses are restored at 12 months (with or without mHTT lowering), the changes in individual lipid species which compose each subclasses correlate with behavioral degradation.

Not all lipid changes were reversed with m*Htt* lowering. Consistent with metabolic defects in HD, here we report reduced levels of species of TG, glycerolipid molecules used for energy storage which can be metabolized by mitochondria, in LacQ140 compared to WT. Of note, reductions in species of TG were not reversed by m*Htt* lowering. Interestingly, the LacQ140 mice exhibited reciprocal changes in the two biosynthetic enzymes *Dgat1* and *Dgat2* at the transcriptional level, suggesting the ability to store energy may arise in part from this variation.

## CONCLUSIONS

Collectively, our studies advocate early lowering of m*HTT* for greatest benefit, and in this context, modest lowering is sufficient to delay some protein and lipid changes. Furthermore, our work shows readily detectable but transient changes in lipids highly enriched in myelin, consistent with possible white matter damage and regeneration occurring in the LacQ140 mouse model.

## Supporting information

Lipidomics subclass and species data for 6-, 9-, and 12-month-old mice

6-month RNAseq full DEG list, GO analysis, GSEA

12-month RNAseq full DEG list, GO analysis, GSEA

Source data and statistics for main figures

Source data and statistics for supplemental figures

## LIST OF ABBREVIATIONS

HD: Huntington’s disease
HTT: Huntingtin
ASOs: Antisense oligonucleotide
miRNA: MicroRNA
shRNA: Short hairpin RNA
DARPP32: Dopamine And CAMP-Regulated Neuronal Phosphoprotein 32
PDE10A: Phosphodiesterase 10A
SCN4B: Sodium Voltage-Gated Channel Beta Subunit 4
ATP5A: ATP Synthase F1 Subunit Alpha
PIP: Phosphatidylinositol phosphate
IPTG: Isopropyl b-D-1-thiogalactopyranoside
Poly-Q: Polyglutamine
GFAP: Glial fibrillary acidic protein
LC-MS: Liquid chromatography – mass spectrometry
PS: Phosphatidylserine
PI: Phosphatidylinositol
BisMePA: Bis-methyl phosphatidic acid
PKC: Protein kinase C
SM: Sphingomyelin
Cer: Ceramide
MGDG: Monogalactosyldiacylglycerol
Hex1Cer: Monohexosylceramide
GalCer: Galactosylceramide
GlcCer: Glucosylceramide
TG: Triacylglycerol
GO BP: Gene Ontology Biological Processes
GSEA: Gene Set Enrichment Analysis
Ugt8a: UDP-Galactose Ceramide Galactosyltransferase
Fa2h: Fatty acid 2-hydroxylase

## DECLARATIONS

### Ethics approval and consent to participate

Mice were housed at Psychogenics (Paramus, NJ) and all treatments and procedures were conducted with oversight by Psychogenics Institutional Animal Care and Use Committee.

### Consent for publication

Not applicable

### Availability of data and materials

All datasets generated are included in this article and can be found in **Additional Files 1-5.** RNA-sequencing data analyzed is available at Gene Expression Omnibus (GEO) accession number GSE156236.

## Authors’ contributions

KKG, MD, and DM conceived experimental plan and wrote manuscript. KS performed all lipidomics and bioinformatics and wrote manuscript, ES performed protein chemistry analysis and wrote manuscript, AB aided in the experimental plan and edited manuscript, CS and SL performed subcellular fractionations.

## Acknowledgements

The authors would like to acknowledge the Dake family for their support.

## MAIN FIGURE LEGENDS

**Supplementary Figure 1.**
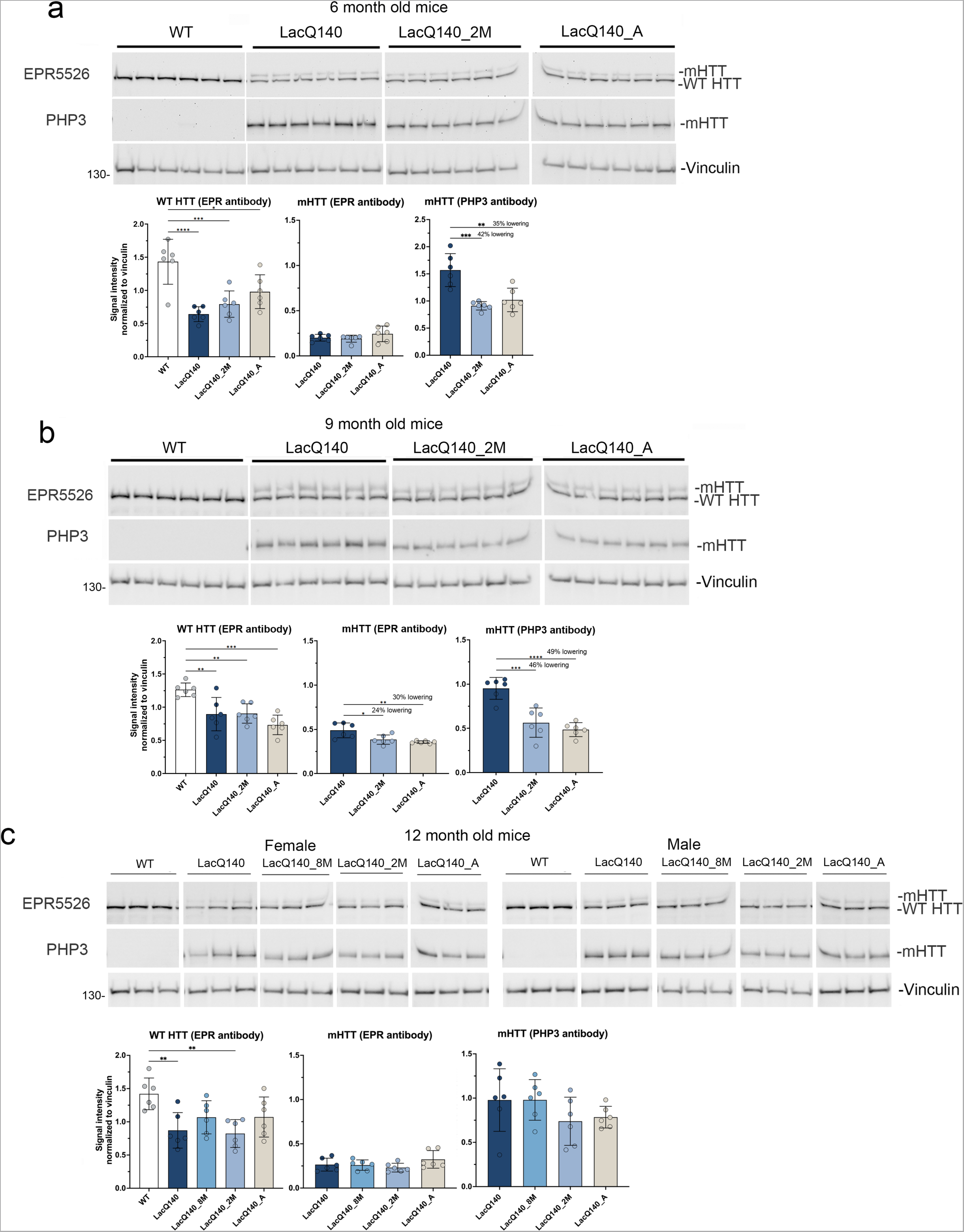
Analysis of mHTT protein levels with EPR5526 and PHP3 in crude homogenates of 6-, 9- and 12-months old mice. HTT levels were analyzed by western blot on equal amounts of protein (10 µg) using anti-HTT antibody EPR5526 and anti-polyQ antibody PHP3. Total pixel intensity quantification for each band measured using ImageJ software in 6- month-old mice shows a significant decrease in WT HTT as detected with EPR5526 in all LacQ140 mice compared to WT mice but no change in mHTT levels (**a**). mHTT levels are significantly lower in LacQ140_2M and LacQ140_A mice as detected with PHP3 compared to LacQ140 (**a**, -42% and -35% respectively, **p<0.01, ***p<0.001, One-way ANOVA with Tukey’s multiple comparison test, n=6). Total pixel intensity quantification in 9-month-old mice shows a significant decrease in WT HTT as detected with EPR5526 in all LacQ140 mice compared to WT mice. (**b**). mHTT levels are significantly lower in LacQ140_2M and LacQ140_A mice as detected with EPR5526 and PHP3 compared to LacQ140 (**b**, EPR5526: -24% and -30%, respectively; PHP3: -46% and -49% respectively, *p<0.05, **p<0.01, ***p<0.001, One-way ANOVA with Tukey’s multiple comparison test, n=6).Total pixel intensity quantification in 12- month-old mice shows a significant decrease in WT HTT as detected with EPR5526 in LacQ140 and LacQ140_2M compared to WT mice. No changes in mHTT levels as detected with either EPR5526 or PHP3 were observed (**c**).

**Supplementary Figure 2.**
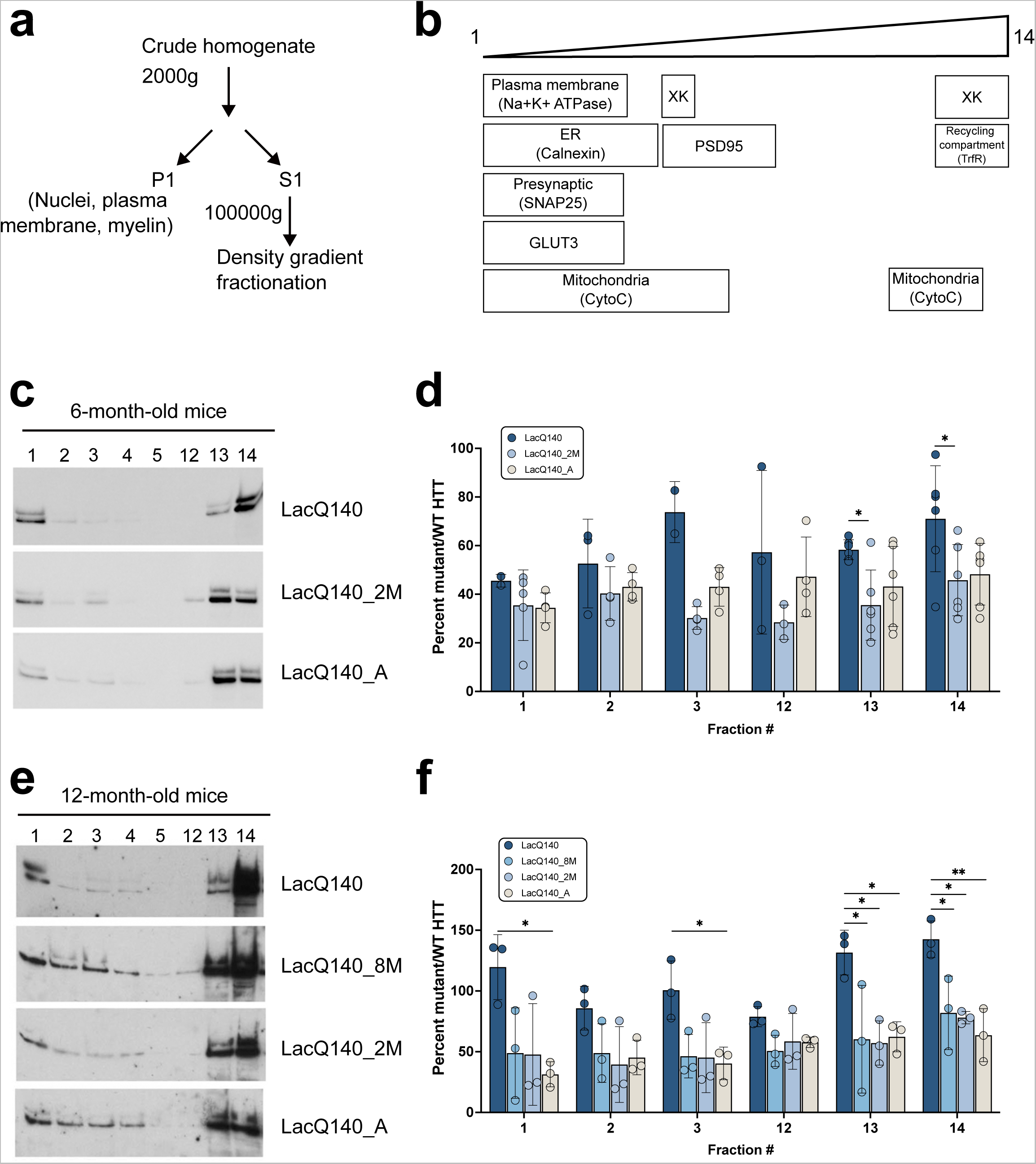
Effects of m*Htt* lowering on the subcellular distribution of WT and mHTT protein by density gradient ultracentrifugation. Diagram depicts the centrifugation strategy for protein samples **(a)**. Schematic shows the approximate location in the fractions of protein markers and the organelles found in these compartments **(b)**. Representative western blot images for equal volumes of fractions 1-5 and 12-14 from 6-month-old mice probed with anti-HTT antibody Ab1 are shown in **(c)**. The remaining images are shown in **Supplementary** Figure 3a. Total pixel intensity quantification for each band measured using ImageJ software is graphed as average percent mutant/WT HTT ± SD for each fraction **(d)**. Since each fraction contains different levels of proteins normally used to control for protein loading, levels of mHTT were normalized to levels of WT HTT which was not repressed/lowered. The ratio mutant/WT HTT is significantly higher in LacQ140 mice compared to LacQ140_2M in fractions 13 and 14 (*p<0.05, One-way ANOVA with Tukey’s multiple comparison test for each fraction, n=6). Representative western blot images for equal volumes of fractions 1-4 and 11-14 from 12-month-old mice probed with anti-HTT antibody Ab1 are shown in **(e)**. The remaining images are shown in **Supplementary** Figure 3c. Total pixel intensity quantification for each band is graphed as average percent mutant/WT HTT ± SD for each fraction **(f)**. The ratio mutant/WT HTT is significantly higher in LacQ140 compared to LacQ140_8M, LacQ140_2M and/or LacQ140_A mice in fractions 1, 3, 13, and 14 (*p<0.05, **p<0.01, One-way ANOVA with Tukey’s multiple comparison test for each fraction, n=3). Graphs indicate data in fractions where mHTT was detected in at least 3 mice except LacQ140_A fractions 1 and 3 where only 2 mice had detectible mHTT.

**Supplementary Figure 3.**
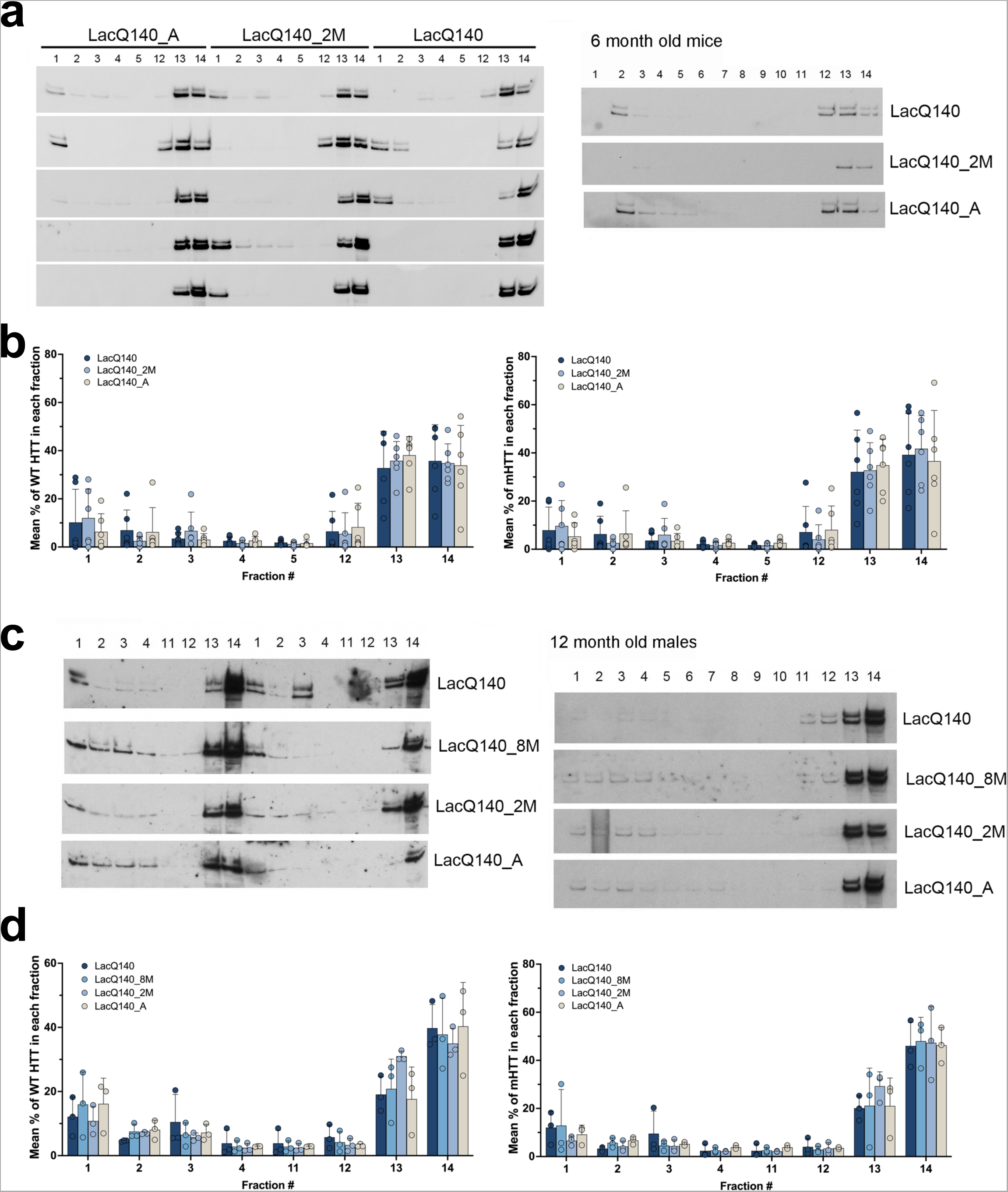
Effects of m*Htt* lowering on the subcellular distribution of WT and mHTT. Western blot images for equal volumes of fractions 1-14 (right blots, each strip is from one mouse) or fractions 1-5 and 12-14 (left blots) from 6-month-old mice probed with anti-HTT antibody Ab1 are shown in **(a)**. Each strip in left blots is a set of 1 mouse per treatment group with the groups labeled at the top of the blots. Total pixel intensity quantification for each band using ImageJ software is graphed as average percent of total HTT signal for each fraction ± SD **(b)**. Representative western blot images for equal volumes of fractions 1-14 (right blots, each strip is from one mouse) or 1-4 and 11-14 (left blots) from 12-month-old mice probed with anti-HTT antibody Ab1 are shown in **(c)**. Each strip in the left blots is from 2 mice per group with the groups labeled on the right. Total pixel intensity quantified for each band using ImageJ software are graphed as average percent of total HTT signal for each fraction ± SD **(d)**. There is no difference in WT or mHTT levels as a percent of total HTT in any fraction with any treatment of LacQ140 mice at 6 or 12 months (One-way ANOVA with Tukey’s multiple comparison test for each fraction, n=6 for 6 months and n=3 for 12 months).

**Supplementary Figure 4.**
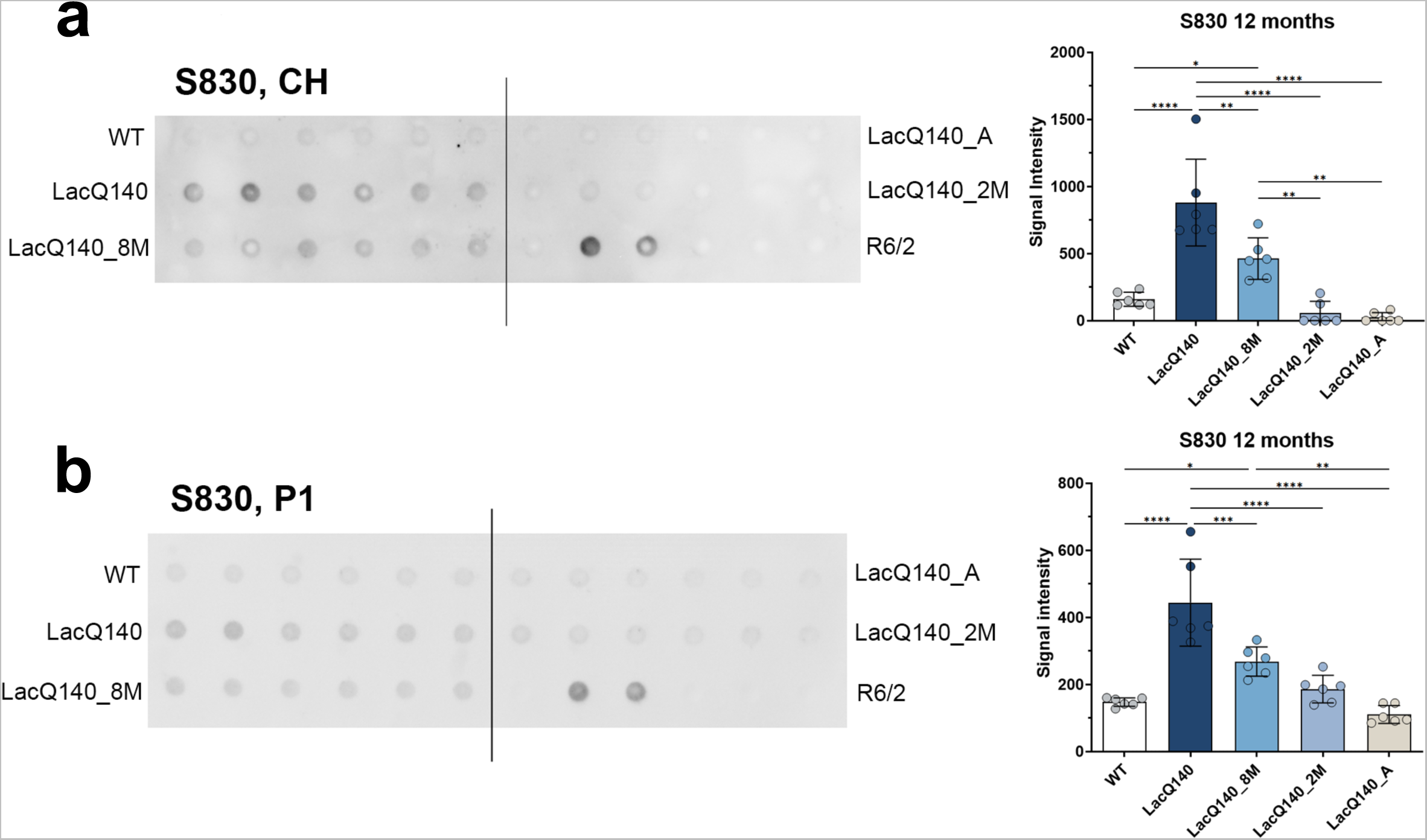
HTT levels in crude homogenates and P1 fractions from WT and LacQ140 mice by filter trap assay. Filter trap assays of 12-month-old crude homogenates **(a)** and P1 fractions **(b)** were probed with S830 antibody. Each dot represents one animal and each of the 6 dots across equals one group which is labeled on the left and right sides. There are 2 dots for the lysates from R6/2 HD mice which have a highly expressing transgene for a small fragment of HTT containing a large CAG repeat (180CAGs) and that accumulate numerous aggregates which have been shown to be retained in the assay and were used as a positive control. There was significantly more signal for aggregated mHTT in the 12-month-old LacQ140 mice compared to WT, LacQ140_8M, LacQ140_2M and LacQ140_A mice (**a**, *p<0.05, **p<0.01, ****p<0.0001, One-way ANOVA with Tukey’s multiple comparison test, n=6). In the P1 fractions, there was significantly more signal for aggregated mHTT detected with S830 antibody in the 12-month LacQ140 mice compared to WT, LacQ140_8M, LacQ140_2M and LacQ140_A mice and in LacQ140_8M compared to WT and LacQ140_A mice (**b**, *p<0.05, **p<0.01, ***p<0.001, ****p<0.0001, One-way ANOVA with Tukey’s multiple comparison test, n=6).

**Supplementary Figure 5.**
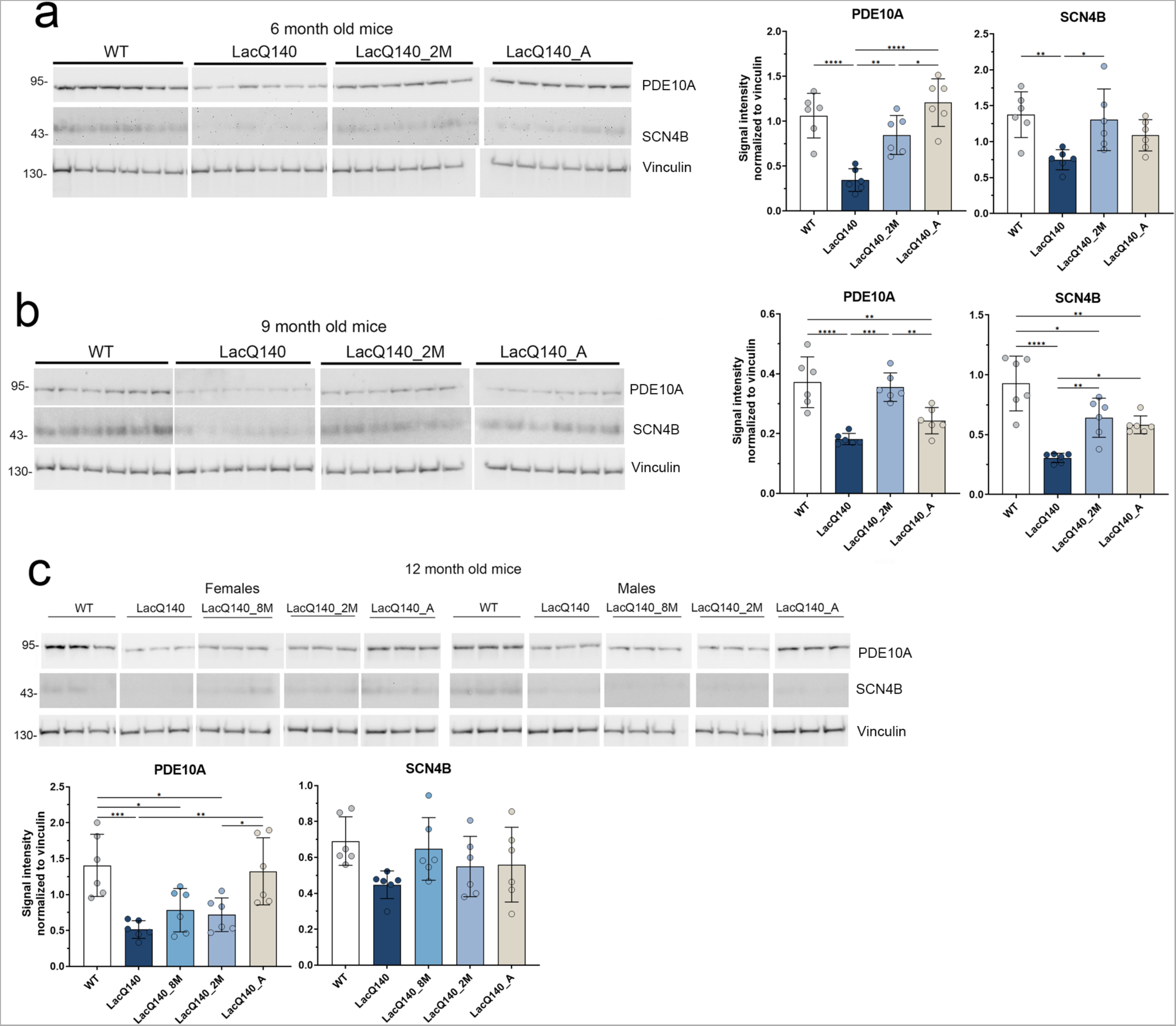
Duration of m*Htt* lowering in 6, 9 and 12 months old LacQ140 mice affects levels of PDE10A and SCN4B. PDE10A and SCN4B levels were analyzed by western blot on equal amounts of protein (10 µg). Total pixel intensity quantification for each band using ImageJ software in 6-month-old mice shows a significant decrease in PDE10A levels in LacQ140 compared to WT mice. There is an increase in PDE10A levels in LacQ140_2M and LacQ140_A mice compared to LacQ140 and no change from WT mice (**a,** *p<0.05, **p<0.01, ****p<0.0001, One-way ANOVA with Tukey’s multiple comparison test, n=6). There is a significant decrease in SCN4B levels in LacQ140 compared to WT mice and a significant increase back to WT levels in LacQ140_2M mice. Total pixel intensity quantification in 9- month-old mice shows a significant decrease in PDE10A levels in LacQ140 compared to WT mice. There is an increase in PDE10A levels in LacQ140_2M compared to LacQ140 and no change from WT mice (**b,** **p<0.01, ***p<0.001, One-way ANOVA with Tukey’s multiple comparison test, n=6). There is a significant decrease in SCN4B levels in LacQ140 compared to WT mice. There is a significant increase in SCN4B levels in LacQ140_2M and LacQ140_A mice compared to LacQ140 but significantly lower than in the WT mice (**b,** *p<0.05, **p<0.01, ***p<0.001, ****p<0.0001, One-way ANOVA with Tukey’s multiple comparison test, n=6). Total pixel intensity quantification in 12-month-old mice shows a significant decrease in PDE10A levels in LacQ140, LacQ140_8M and LacQ140_2M compared to WT mice. There is an increase in PDE10A levels in LacQ140_A mice compared to LacQ140 and LacQ140_2M and no change from WT mice (**c,** *p<0.05, **p<0.01, ***p<0.001, One-way ANOVA with Tukey’s multiple comparison test, n=6). There are no changes in SCN4B levels in any of the LacQ140 or WT mice.

**Supplementary Figure 6.**
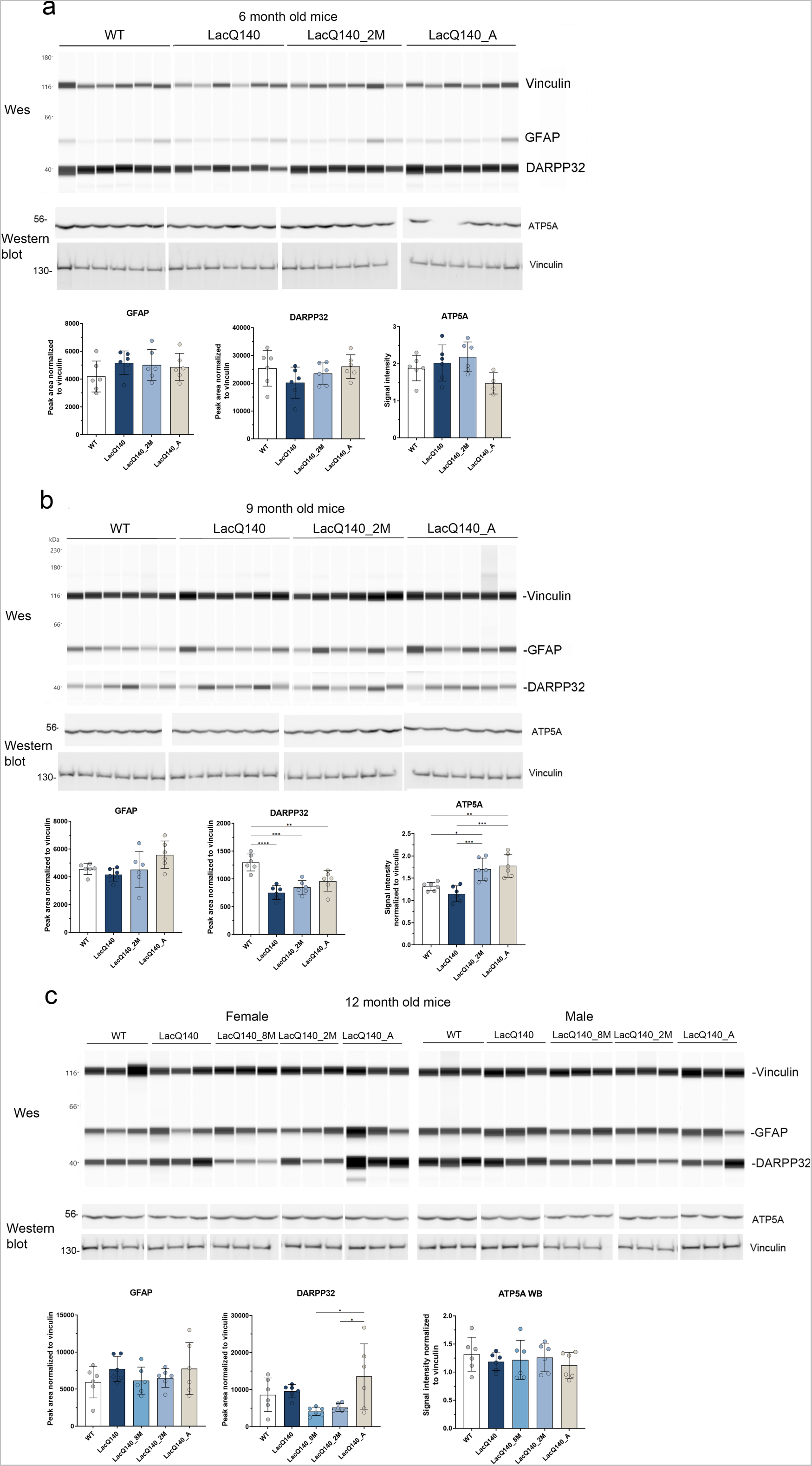
Duration of m*Htt* lowering in 6, 9 and 12 months old LacQ140 mice has minimal effects on levels of GFAP, DARPP32 and ATP5A. GFAP and DARPP32 levels were analyzed by capillary immunoassay on equal amounts of protein (0.6 µg) and ATP5A levels were analyzed by western blot on equal amounts of protein (10 µg). In 6-month-old mice, peak area analysis shows no significant change in GFAP or DARPP32 levels (**a**) and total pixel intensity quantification shows no changes in ATP5A levels in any of the LacQ140 or WT mice (**a**). Peak area analysis in 9-month-old mice shows a significant decrease in DARPP32 levels in all LacQ140 mice compared to WT mice (**b**, **p<0.01, ***p<0.001, One-way ANOVA with Tukey’s multiple comparison test, n=6). Total pixel intensity quantification shows ATP5A levels are significantly higher in LacQ140_2M and LacQ140_A mice compared to WT or LacQ140 mice (**b,** *p<0.05, **p<0.01, ***p<0.001, One-way ANOVA with Tukey’s multiple comparison test, n=6). Peak area analysis in 12-month-old mice shows no significant change in GFAP levels in LacQ140 or WT mice. There is a significant decrease in DARPP32 levels in LacQ140_8M and LacQ140_2M compared to LacQ140_A mice (**c,** *p<0.05, One-way ANOVA with Tukey’s multiple comparison test, n=6). Total pixel intensity quantification for each band using ImageJ software shows no changes in ATP5A levels in any of the LacQ140 or WT mice.

**Supplementary Figure 7.**
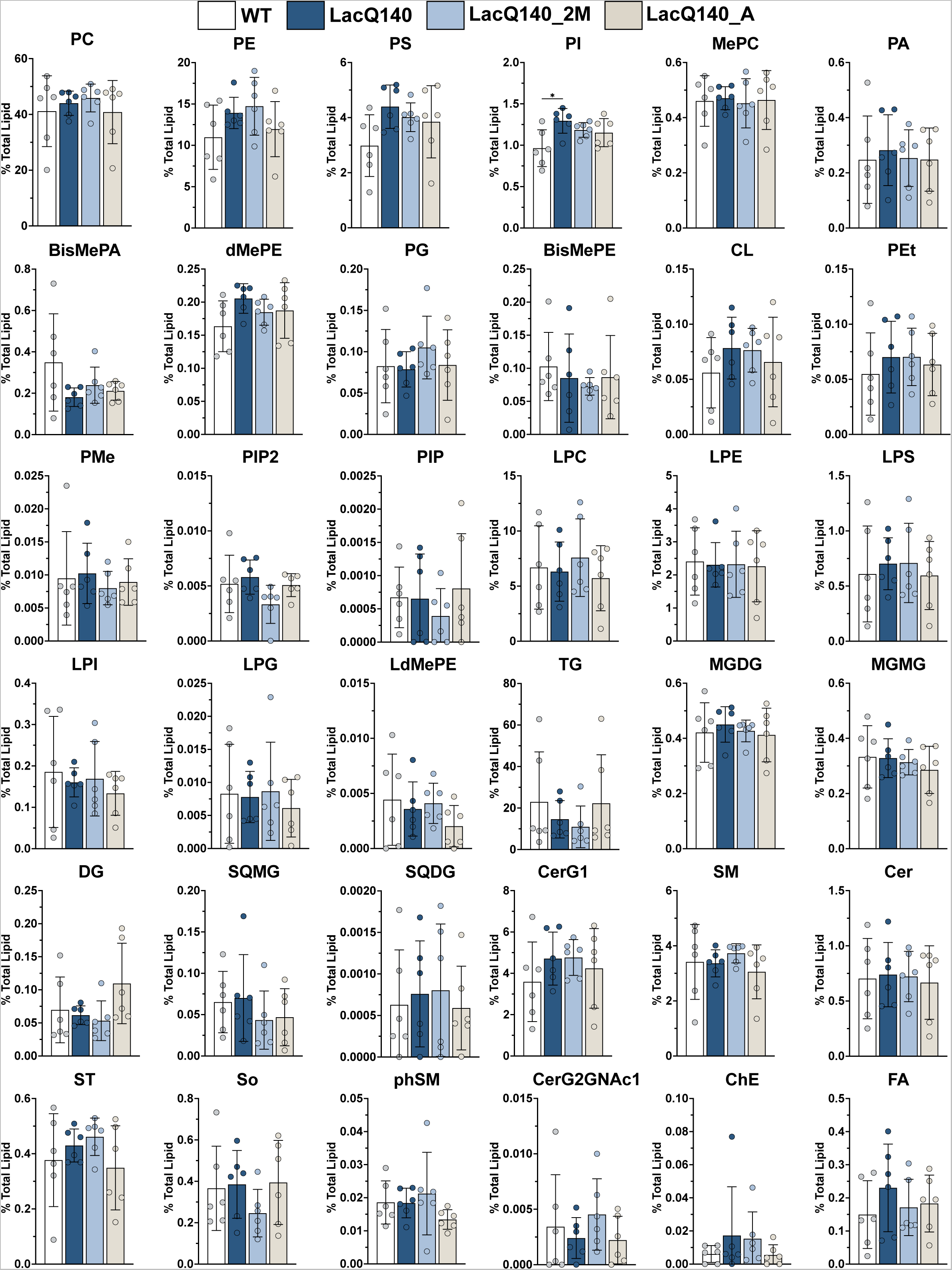
Lipid subclasses detected in caudate putamen of 6-month-old mice. Lipids were extracted and analyzed by LC-MS/MS. Graphs show relative intensities for indicated lipid subclasses expressed as a percent of total lipid intensity per sample for each genotype or treatment group. Plotted values represent summed lipid subclass intensity standardized to total amount of lipid detected in the same sample. Bar charts underneath individual points represent group means and error bars are ± standard deviation. PI was significantly increased in striatum of 6-month-old LacQ140 mice compared to WT mice (One-way ANOVA with Tukey’s multiple comparisons test, F(3, 20) = 4.176, * P=0.0189, n=6). This increase is not significant when p-values are adjusted for multiple testing using the Benjamini, Krieger and Yekutieli procedure with a false discovery rate of 5%, (q = 0.7144, N=36 subclasses). No changes between groups were found in any other subclasses. Abbreviations: *<u>Glycerophospholipids</u>:* PC = phosphatidylcholine, PE = phosphatidylethanolamine, PS = phosphatidylserine, PI = phosphatidylinositol, MePC = methylphosphocholine, PA = phosphatidic acid, BisMePA = bis-methyl phosphatidic acid, dMePE = dimethyl phosphatidylethanolamine, PG = phosphatidylgylcerol, BisMePE = bis-methyl phosphatidyl ethanolamine, CL = cardiolipin, PEt = phosphatidylethanol, PMe = phosphatidylmethanol, PIP2 = phosphatidylinositol-bisphosphate, PIP = phosphatidylinositol-monophosphate, LPC = lysophosphatidylcholine, LPE = lysophosphatidylethanolamine, LPS = lysophosphatidylserine, LPI = lysophosphatidylinositol LPG = lysophosphosphatidylgylcerol, LdMePE = lysodimethyl phosphatidyl ethanolamine *<u>Glycerolipids:</u>* TG = triacylglycerol, MGDG = monogalactosyldiacylglycerol, MGMG = monogalactosylmonoacylglycerol, DG = diacylglycerol, SQMG = sulfoquinovosylmonoacylglycerol, SQDG = sulfoquinovosyldiacylglycerol, *<u>Sphingolipids</u>:* CerG1 = simple Glc series, SM = sphingomyelin, Cer = ceramide, ST = sulfatide, So = sphingosine, phSM = sphingomyelin phytosphingosine, CerG2GNAc1 = Simple Glc series *<u>Sterol lipids:</u>* ChE = cholesterol ester *<u>Fatty acyls:</u>* FA = fatty acid

**Supplementary Figure 8.**
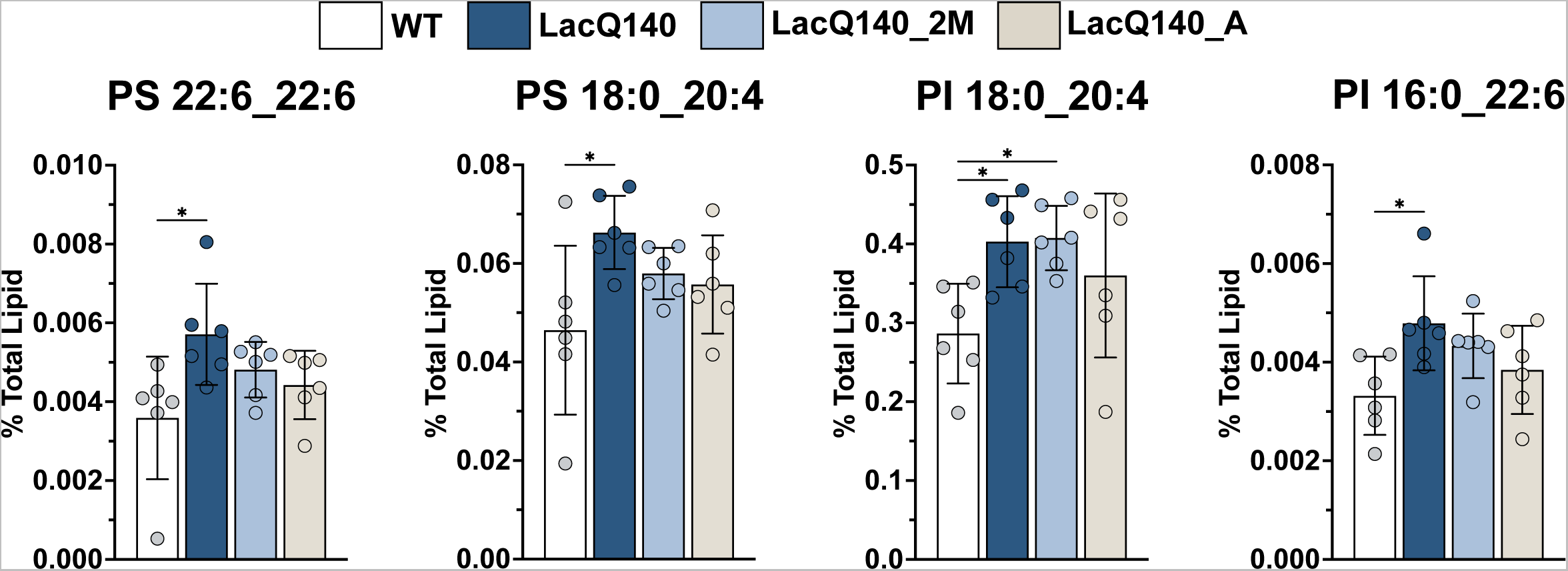
Individual lipid species significantly different between 6-month-old LacQ140 and WT mice. Lipids were extracted and analyzed by LC-MS/MS. Graphs show relative intensities for indicated lipid species expressed as a percent of total lipid intensity per sample for each genotype or treatment group. Plotted values represent individual lipid species intensity standardized to total amount of lipid detected in the same sample. Bar charts underneath individual points represent group means and error bars are ± standard deviation. One-way analysis of variance (ANOVA) was used to evaluate differences in lipid species intensity between groups and Tukey’s multiple comparisons test was used for post-hoc pairwise comparisons (n=6 mice per group, *p < 0.05, Tukey’s multiple comparisons test). To correct for multiple testing over all lipids analyzed, the Benjamini, Krieger and Yekutieli procedure was used with a false discovery rate of 5% (N=800 lipid species), FDR adjusted p-values are reported as q values. No individual species found to be different by one-way ANOVA were significant following correction of p-values. PS 22:6_22:6: F(3, 20) = 3.812,*P=0.0260, q=1, ns; PS 18:0_20:4: F(3, 20) = 3.349, *P=0.0396, q=1, ns; PI 18:0_20:4: F(3, 20) = 3.812, *P=0.0260, q=1, ns; PI 16:0_22:6: F(3, 20) = 3.466, *P=0.0356, q=1, ns. Abbreviations: PI = phosphatidylinositol, PS = phosphatidylserine

**Supplementary Figure 9.**
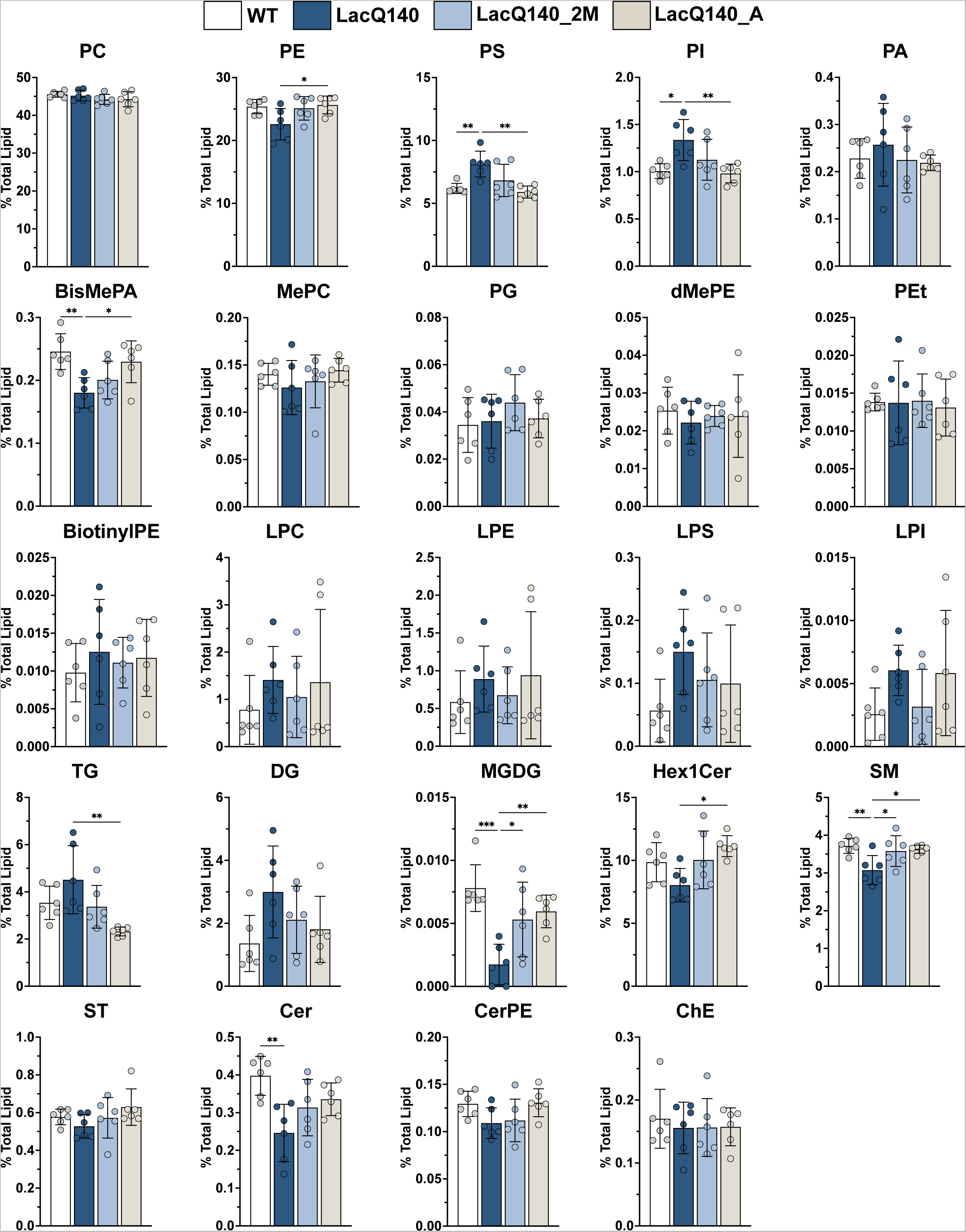
Lipid subclasses detected in caudate putamen of 9-month-old mice. Lipids were extracted and analyzed by LC-MS/MS. Graphs show relative intensities for indicated lipid subclasses expressed as a percent of total lipid intensity per sample for each genotype or treatment group. Plotted values represent summed lipid subclass intensity standardized to total amount of lipid detected in the same sample. Bar charts underneath individual points represent group means and error bars are ± standard deviation. One-way analysis of variance (ANOVA) was used to evaluate differences in lipid subclass intensity between groups and Tukey’s multiple comparisons test was used for post-hoc pairwise comparisons (n=6 mice per group, *p < 0.05, **p < 0.01, Tukey’s multiple comparisons test). To correct for multiple testing the Benjamini, Krieger and Yekutieli procedure was used with a false discovery rate of 5% (N=24 subclasses) and FDR adjusted p-values are reported as q values. PS: F(3, 20) = 7.601, **P=0.0014, q=0.0125, PI: F(3, 20) = 5.707, **P=0.0054, q=0.0168, BisMePA: F(3, 20) = 6.086, **P=0.0041,q=0.0168, SM: F(3, 20) = 5.465, **P=0.0066, q=0.0168, Cer: F(3, 20) = 5.883, **P=0.0048, q=0.0168, MGDG: F(3, 20) = 9.350, ***P=0.0005, q=0.0089. PE, TG and Hex1Cer were unchanged between LacQ140 and WT groups but had changes among other treatment groups (PE: F(3, 20) = 3.717, *P=0.0284, q=0.0563, ns, TG: F(3, 20) = 5.637, **P=0.0057, q=0.0168, Hex1Cer: F(3, 20) = 3.891, *P=0.0243, q=0.054, ns). Abbreviations: *<u>Glycerophospholipids:</u>* PC = phosphatidylcholine, PE = phosphatidylethanolamine, PS = phosphatidylserine, PI = phosphatidylinositol, PA = phosphatidic acid, BisMePA = bis-methyl phosphatidic acid, MePC = methylphosphocholine, PG = phosphatidylgylcerol, dMePE = dimethyl phosphatidylethanolamine, PEt = phosphatidylethanol, BiotinylPE = biotinyl-phosphoethanolamine, LPC = lysophosphatidylcholine, LPE = lysophosphatidylethanolamine, LPS = lysophosphatidylserine, LPI = lysophosphatidylinositol *<u>Glycerolipids:</u>* TG = triacylglycerol, DG = diacylglycerol, MGDG = monogalactosyldiacylglycerol *<u>Sphingolipids:</u>* Hex1Cer = monohexosylceramide, SM = sphingomyelin, ST = sulfatide, Cer = ceramide, CerPE = ceramide phosphorylethanolamine *<u>Sterol lipids</u>*: ChE = cholesterol ester

**Supplementary Figure 10.**
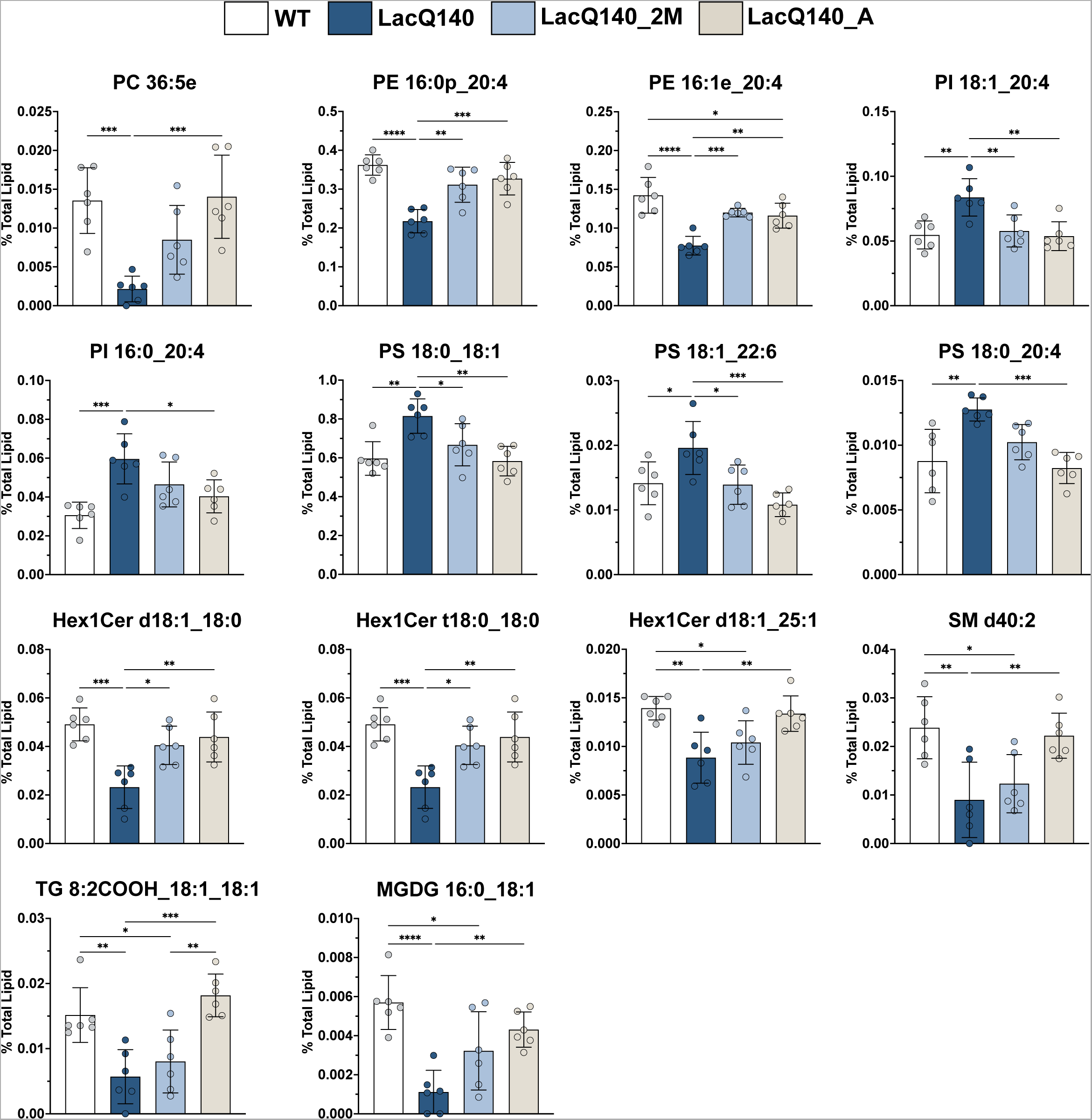
Individual lipid species significantly changed between 9-month-old LacQ140 and WT mice. Lipids were extracted and analyzed by LC-MS/MS. Graphs show relative intensities for indicated lipid species expressed as a percent of total lipid intensity per sample for each genotype or treatment group. Plotted values represent individual lipid species intensity standardized to total amount of lipid detected in the same sample. Bar charts underneath individual points represent group means and error bars are ± standard deviation. One-way analysis of variance (ANOVA) was used to evaluate differences in lipid species intensity between groups and Tukey’s multiple comparisons test was used for post-hoc pairwise comparisons (n=6 mice per group, *p < 0.05, **p < 0.01, ***p < 0.001, ****p < 0.0001, Tukey’s multiple comparisons test). To correct for multiple testing over all lipids analyzed, the Benjamini, Krieger and Yekutieli procedure was used with a false discovery rate of 5% (N=632 lipid species), FDR adjusted p-values are reported as q values. PC 36:5e: F(3, 20) = 10.68, ***P=0.0002, q=0.02, PE 16:1e_20:4: F(3, 20) = 18.14, ****P<0.0001, q=0.003, PE 16:0p_20:4: F(3, 20) = 16.97, ****P<0.0001, q=0.003, PI 18:1_20:4: F(3, 20) = 8.086, **P=0.0010, q=0.041, PI 16:0_20:4: F(3, 20) = 8.449, ***P=0.0008, q=0.039, PS 18:0_18:1: F(3, 20) = 8.226, ***P=0.0009, q=0.041, PS 18:1_22:6: F(3, 20) = 7.916, **P=0.0011, q=0.042, PS 18:0_20:4: F(3, 20) = 9.739, ***P=0.0004, q=0.023, Hex1Cer d18:1_18:0: F(3, 20) = 10.37, ***P=0.0002, q=0.02, Hex1Cer t18:0_18:0: F(3, 20) = 10.35, ***P=0.0003, q=0.02, Hex1Cer d18:1_25:1: F(3, 20) = 8.440, ***P=0.0008, q=0.039, SM d40:2: F(3, 20) = 8.052, **P=0.0010, q=0.041, TG 8:2COOH_18:1_18:1: F(3, 20) = 11.98, ***P=0.0001, q=0.017, MGDG 16:0_18:1: F(3, 20) = 11.28, ***P=0.0002, q=0.019. Abbreviations: *<u>Glycerophospholipids:</u>* PC = phosphatidylcholine, PE = phosphatidylethanolamine, PI = phosphatidylinositol, PS = phosphatidylserine *<u>Sphingolipids:</u>* Hex1Cer = monohexosylceramide, SM = sphingomyelin *<u>Glycerolipids:</u>* MGDG = monogalactosyldiacylglycerol, TG = triacylglycerol

**Supplementary Figure 11.**
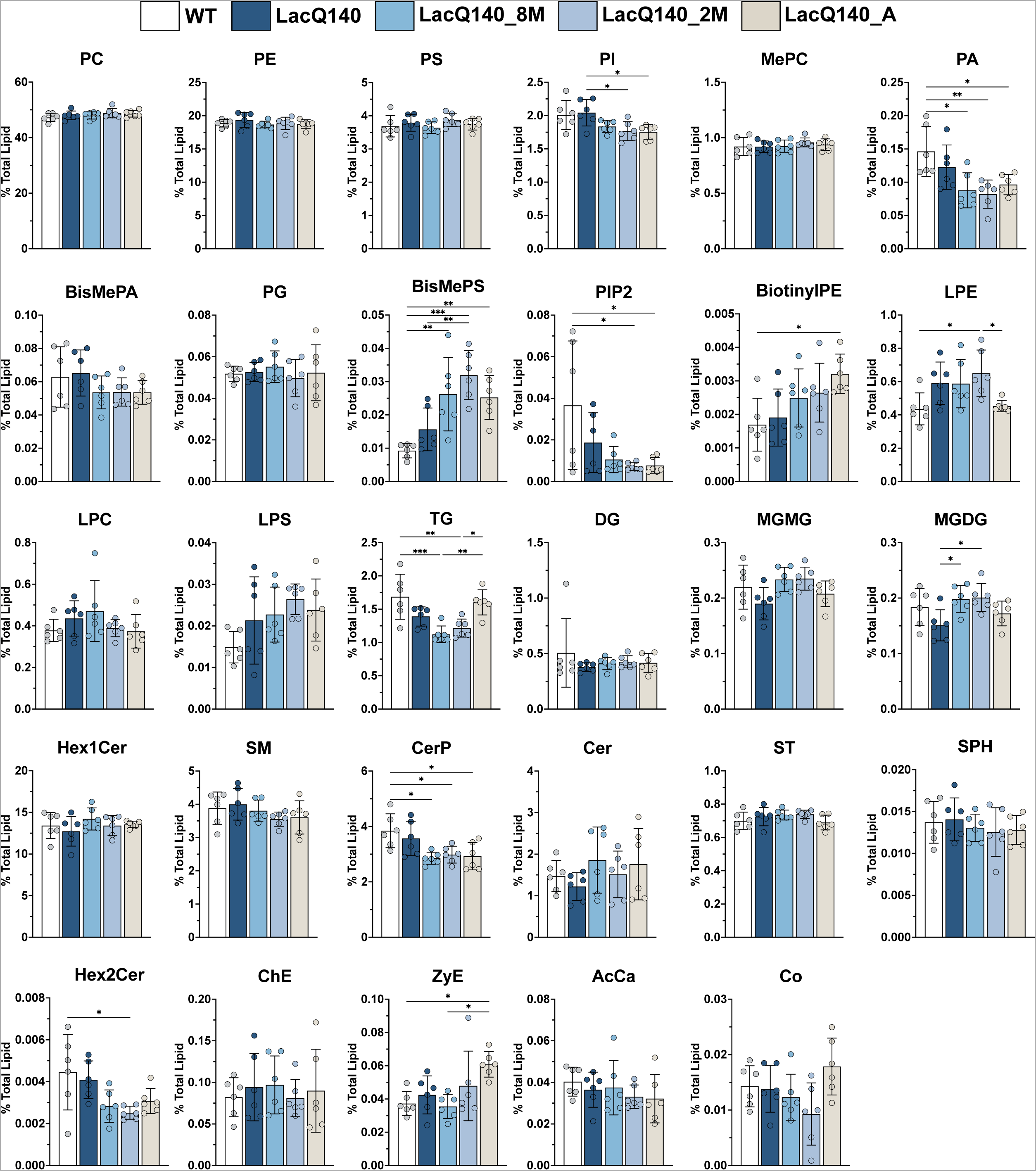
Lipid subclasses detected in caudate putamen of 12-month-old mice. Lipids were extracted and analyzed by LC-MS/MS. Graphs show relative intensities for indicated lipid subclasses expressed as a percent of total lipid intensity per sample for each genotype or treatment group. Plotted values represent summed lipid subclass intensity standardized to total amount of lipid detected in the same sample. Bar charts underneath individual points represent group means and error bars are ± standard deviation. One-way analysis of variance (ANOVA) was used to evaluate differences in lipid subclass intensity between groups and Tukey’s multiple comparisons test was used for post-hoc pairwise comparisons (n=6 mice per group, *p < 0.05, **p < 0.01, ***p < 0.001, Tukey’s multiple comparisons test). To correct for multiple testing the Benjamini, Krieger and Yekutieli procedure was used with a false discovery rate of 5% (N=29 subclasses), FDR adjusted p-values are reported as q values. PI: F(4, 25) = 4.316, **P=0.0086, q=0.0338, PA: F(4, 25) = 5.537, **P=0.0025, q=0.0184, BisMePS: F(4, 25) = 9.308, ****P<0.0001, q=0.0022, PIP2: F(4, 25) = 3.743, *P=0.0161, q=0.0394, BiotinylPE: F(4, 25) = 3.446, *P=0.0225, q=0.0451, LPE: F(4, 25) = 3.957, *P=0.0127, q=0.035. TG: F(4, 25) = 8.593, ***P=0.0002, q=0.0022, MGMG: F(4, 25) = 2.765, *P=0.0497, q=0.0887, ns, MGDG: F(4, 25) = 3.491, *P=0.0214, q=0.0451. CerP: F(4, 25) = 5.185, **P=0.0035, q=0.0193, Hex2Cer: F(4, 25) = 4.076, *P=0.0112, q=0.035, ZyE: F(4, 25) = 4.256, **P=0.0092, q= 0.0338. Abbreviations: *<u>Glycerophospholipids:</u>* PC = phosphatidylcholine, PE = phosphatidylethanolamine, PS = phosphatidylserine, PI = phosphatidylinositol, MePC = methylphosphocholine, PA = phosphatidic acid, BisMePA = bis-methyl phosphatidic acid, PG = phosphatidylgylcerol, BisMePS = bis-methylphosphatidylserine, PIP2 = phosphatidylinositol-bisphosphate, BiotinylPE = biotinyl-phosphoethanolamine, LPE = lysophosphatidylethanolamine, LPC = lysophosphatidylcholine, LPS = lysophosphatidylserine *<u>Glycerolipids:</u>* TG = triacylglycerol, DG = diacylglycerol, MGMG = monogalactosylmonoacylglycerol, MGDG = monogalactosyldiacylglycerol *<u>Sphingolipids:</u>* Hex1Cer = monohexosylceramide, SM = sphingomyelin, CerP = ceramide phosphate, Cer = ceramide, ST = sulfatide, SPH = sphingoid base, Hex2Cer = dihexosylceramide <u>Sterol lipids:</u> ChE = cholesterol ester, ZyE = zymosterol *<u>Fatty acyls:</u>* AcCa = acyl carnitine *<u>Prenol lipids:</u>* Co = coenzyme

**Supplementary Figure 12.**
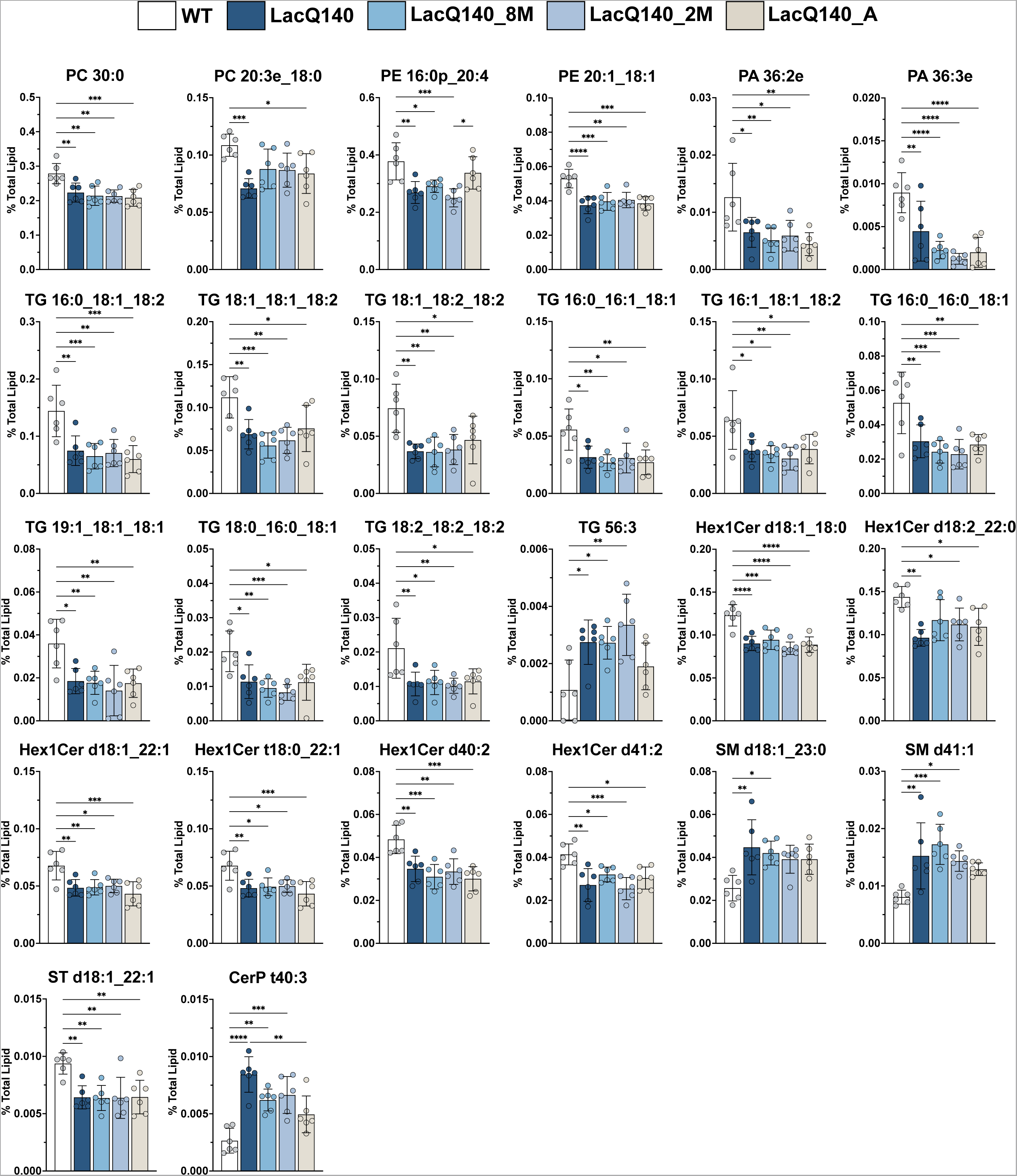
Individual lipid species significantly changed between 12-month-old LacQ140 and WT mice. Lipids were extracted and analyzed by LC-MS/MS. Graphs show relative intensities for indicated lipid species expressed as a percent of total lipid intensity per sample for each genotype or treatment group. Plotted values represent individual lipid species intensity standardized to total amount of lipid detected in the same sample. Bar charts underneath individual points represent group means and error bars are ± standard deviation. One-way analysis of variance (ANOVA) was used to evaluate differences in lipid species intensity between groups and Tukey’s multiple comparisons test was used for post-hoc pairwise comparisons (n=6 mice per group, *p < 0.05, **p < 0.01, ***p < 0.001, ****p < 0.0001, Tukey’s multiple comparisons test). To correct for multiple testing over all lipids analyzed, the Benjamini, Krieger and Yekutieli procedure was used with a false discovery rate of 5% (N=735 lipid species), FDR adjusted p-values are reported as q values. PC 30:0: F(4, 25) = 7.601, ***P=0.0004, q=0.0132, PC 20:3e_18:0: F(4, 25) = 5.567, **P=0.0024, q=0.0327, PE 16:0p_20:4: F(4, 25) = 7.879, ***P=0.0003, q=0.0114, PE 20:1_18:1: F(4, 25) = 10.36, ****P<0.0001, q=0.0038, PA 36:2e: F(4, 25) = 5.630, **P=0.0023, q=0.0327, PA 36:3e: F(4, 25) = 13.27, ****P<0.0001, q=0.0007, TG 16:0_18:1_18:2: F(4, 25) = 8.357, ***P=0.0002, q=0.0094, TG 18:1_18:1_18:2: F(4, 25) = 7.113, ***P=0.0006, q=0.0182, TG 18:1_18:2_18:2: F(4, 25) = 6.249, **P=0.0013, q=0.0272, TG 16:0_16:1_18:1: F(4, 25) = 5.794, **P=0.0019, q=0.0306, TG 16:1_18:1_18:2: F(4, 25) = 4.983, **P=0.0043, q=0.0438, TG 16:0_16:0_18:1: F(4, 25) = 7.833, ***P=0.0003, q=0.0114, TG 19:1_18:1_18:1: F(4, 25) = 6.060, **P=0.0015, q=0.0272, TG 18:0_16:0_18:1: F(4, 25) = 6.726, ***P=0.0008, q=0.0217, TG 18:2_18:2_18:2: F(4, 25) = 5.352, **P=0.0030, q=0.0345, TG 56:3: F(4, 25) = 6.109, **P=0.0014, q=0.0272, Hex1Cer d18:1_18:0: F(4, 25) = 14.49, ****P<0.0001, q=0.0005, Hex1Cer d18:2_22:0: F(4, 25) = 5.594, **P=0.0023, q=0.0327, Hex1Cer d18:1_22:1: F(4, 25) = 6.637, ***P=0.0009, q=0.0219, Hex1Cer t18:0_22:1: F(4, 25) = 6.181, **P=0.0013, q=0.0272, Hex1Cer d40:2: F(4, 25) = 9.257, ****P<0.0001, q=0.0058, Hex1Cer d41:2: F(4, 25) = 7.880, ***P=0.0003, q=0.0114, SM d18:1_23:0: F (4, 25) = 5.007, **P=0.0042, q=0.0438, SM d41:1: F(4, 25) = 6.980, ***P=0.0006, q=0.0197, ST d18:1_22:1: F (4, 25) = 6.263, **P=0.0012, q=0.0272, CerP t40:3: F (4, 25) = 14.26, ****P<0.0001, q=0.0005. Abbreviations: *<u>Glycerophospholipids:</u>* PC = phosphatidylcholine, PE = phosphatidylethanolamine, PA = phosphatidic acid *<u>Glycerolipids:</u>* TG = triacylglycerol *<u>Sphingolipids:</u>* Hex1Cer = monohexosylceramide, SM = sphingomyelin, ST = sulfatide, CerP = ceramide phosphate

**Supplementary Figure 13.**
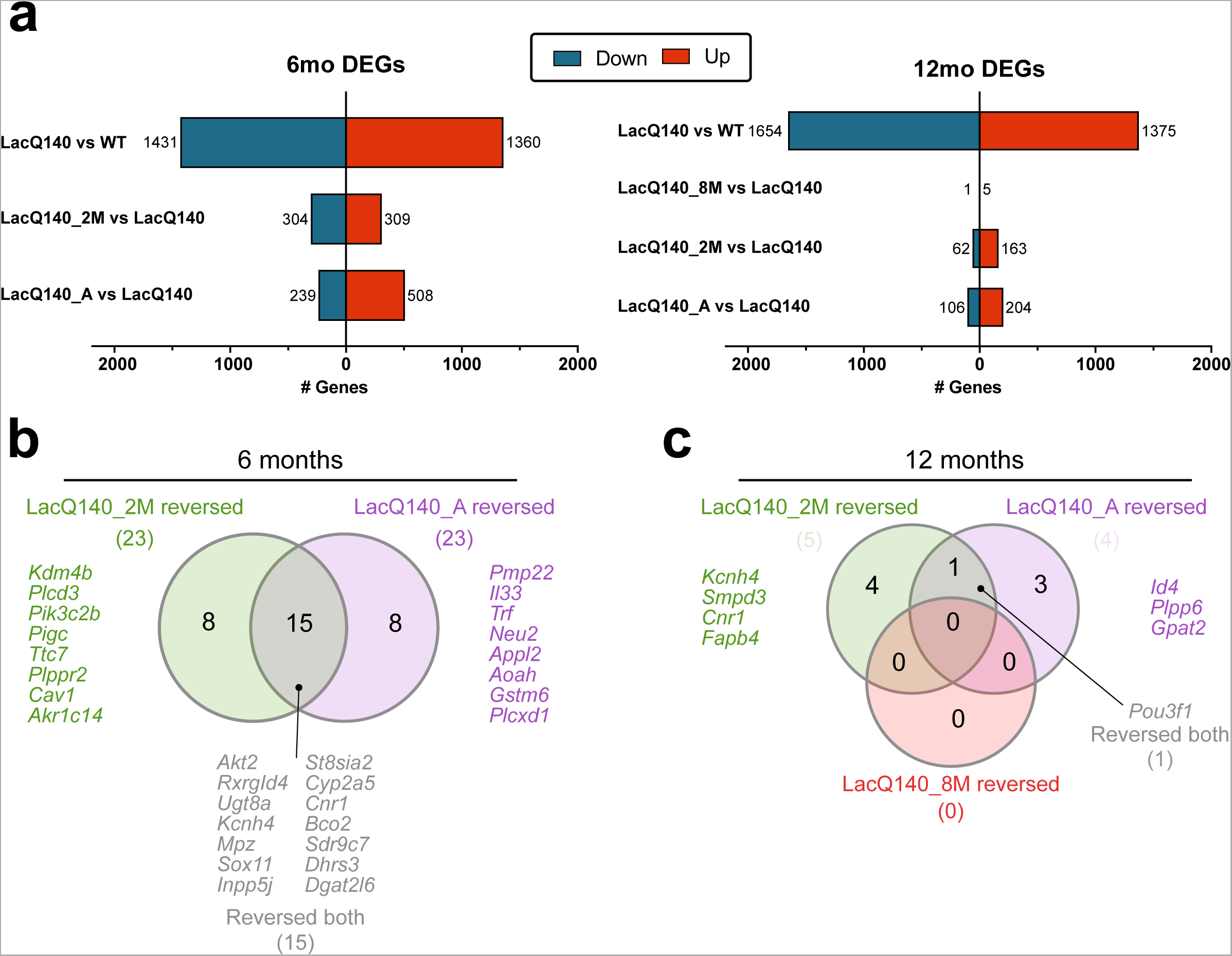
DEGs, reversed genes in 6- and 12-month-old LacQ140 mice. **(a)** Bar chart shows number of differentially expressed genes (DEGs) at 6-months (left) and 12-months (right). Total number of DEGs in LacQ140 mice (upregulated: red and downregulated: blue) are shown in the top bar; lower bars show number of DEGs in mice with mHtt repression (LacQ140_8M, LacQ140_2M, LacQ140_A) compared to LacQ140 mice (i.e., genes “reversed” with m*Htt* repression). **(b)** Venn diagram depicts genes “reversed” at 6-months for each respective m*Htt* repression group as shown in Figure 6c. **(c)** Venn diagram depicts genes “reversed” at 12-months for each respective m*Htt* repression group as shown in Figure 6f.

